# The human claustrum activates across multiple cognitive tasks

**DOI:** 10.1101/2024.11.20.624607

**Authors:** Celine H.L. Huang, Brent W. Stewart, Chun Yin Liu, Phivos Phylactou, Brian N. Mathur, David A. Seminowicz

**Author notes:** Correspondence should be addressed to David A. Seminowicz.

## Abstract

Cognitive control, the ability to manage information during purposeful actions, is crucial for everyday functioning and can become impaired in a variety of neuropsychiatric disorders. The claustrum, a subcortical brain structure, has recently been implicated in functional mechanisms underlying cognitive control. A current theory on the claustrum’s function, the Network Instantiation in Cognitive Control (NICC) model, proposes that the claustrum acts as a cortical network hub synchronizing distant parts of the brain to optimize task performance across cognitive domains. Testing this in this study, we examined the claustrum signal within a dataset (n = 55) that includes functional MRI (fMRI) of healthy participants engaged in four well-established cognitive tasks: the Stroop task, AX-continuous performance task (AX-CPT), cued task-switching, and Sternberg working memory task. Bilateral claustrum activation was observed during certain conditions and trial phases of all four tasks, particularly during active use of cognitive control, and coinciding with task-positive cortical network activations. These findings provide further support for the NICC model of claustrum function, demonstrating claustrum activation across multiple cognitive tasks, and potentially paving the way for new insights into how cognitive processes can become compromised in neuropsychiatric disorders.

## Introduction

Cognitive control, the ability to regulate information processing for goal-oriented behaviour (Mackie et al., 2013), is crucial for survival and is viewed to operate through dynamically organized, large-scale cortical networks (Bressler, 1995; Ji et al., 2019; Thomas Yeo et al., 2011). Among these are task-positive networks, which emerge from the coordinated activation of distributed brain regions during goal-directed task execution (Corbetta & Shulman, 2002; Fox et al., 2005), one example being the frontoparietal network (Dosenbach et al., 2007, 2008; Vincent et al., 2008). Given that cortical network dysfunction is tightly linked to cognitive impairment in neuropsychiatric conditions (Amboni et al., 2015; Dai & He, 2014; Rocca et al., 2018), understanding the circuit mechanisms that govern cortical network control is essential for developing targeted interventions that can improve cognitive function and provide deeper insights into how these networks operate to support healthy brain function.

The claustrum, a thin, sheet-like subcortical structure, has recently been implicated in supporting cortical networks for cognitive control. This structure possesses bidirectional connections with widespread areas of the cerebral cortex and receives input from select subcortical structures (Judith et al., 2002; Milardi et al., 2015; Torgerson et al., 2015; Wang et al., 2017; White et al., 2017). This connectivity enables the claustrum to synaptically link distant cortical regions (Chia et al., 2020; Qadir et al., 2022; White et al., 2018), and is thought to facilitate the co-activation of these regions as coordinated neural networks (Barrett et al., 2020; Qadir et al., 2022; Rodríguez-Vidal et al., 2024). Human fMRI resting state and task-based data support the notion that this synaptic connectivity of the claustrum is integral to cortical network architecture, as the claustrum exhibits functional and effective connectivity with cognitive control-related network regions (Krimmel et al., 2019; Stewart et al., 2024). A recent theory, termed the Network Instantiation in Cognitive Control (NICC) model, outlines the claustrum as a key network hub in the brain that instantiates a cortical network with appropriate network integrity to support cognitive control (Madden et al., 2022). As such, the NICC model predicts that the claustrum activates under a variety of cognitively demanding conditions to instantiate task-positive networks. This prediction has yet to be explicitly tested.

A diverse array of cognitive challenges are known to engage task-positive networks (Cabeza & Nyberg, 2000), each recruiting a unique subset of cognitive functions based on the task’s demands, such as selective attention, working memory, and executive control. The purpose of the current study is to test the NICC model of claustrum function by examining claustrum activation across a variety of cognitive tasks. We hypothesized that claustrum activation would be associated with tasks that activate task-positive networks, specifically during cognitively demanding conditions. We observed significant activation of task-positive networks and bilateral claustrum upon immediate challenge of cognitively demanding task conditions throughout the Stroop, AX-CPT, and Sternberg task. These findings support the NICC model of claustrum function, advancing our understanding of the role of the claustrum and the neurobiological mechanisms underlying cognitive control.

## Materials & Methods

### Participants

This study uses a publicly available dataset, titled the Dual Mechanisms of Cognitive Control or DMCC55B (Etzel et al., 2022), sourced from OpenNeuro (Braver et al., 2021). The dataset contains task-based fMRI and anatomical data from 55 healthy participants (Mean age = 31.7 years, SD = 5.9; 34 female). Inclusion criteria for this study included screening for MRI safety contraindications, an age of between 18 and 45 years of age, without severe mental illness or neurological trauma, and restricted drug/medication usage. All participants completed an out-of-scanner behavioural session consisting of task training with illustrated instructions, practice trials, and scanner familiarization. Full participant selection and screening information can be found at Etzel et al. (2022).

### Imaging Sessions

All scans were acquired using a 3 T Siemens Prisma scanner with a 32-channel head coil in the MR Facility of the Washington University Medical Center. Functional (blood oxygenation level dependent; BOLD) scans were acquired with CMRR multiband sequences, multiband (simultaneous multislice) factor 4, without in-plane acceleration (iPat = none), resulting in 2.4 mm isotropic voxels and a 1.2 s TR (TE 33 msec, flip angle 63°). T1 and T2-weighted high-resolution structural scans were acquired at the beginning of the scanning session (T1: 0.8 mm isotropic voxels; 2.4 s TR, 0.00222 s TE, 1 s TI, 8° flip angle; T2: 0.8 mm isotropic voxels; 3.2 s TR, 0.563 s TE, 120° flip angle). Imaging sessions typically began with the anatomical scans, followed by the first resting state run (5 minutes), the first two tasks (four runs total; two of each task), the second resting state run (also 5 minutes), and finally the remaining two pairs of task runs (also four runs total; two of each task).

### Data Preprocessing

Preprocessing of the fMRI data was performed by Etzel et al. (2022) using fMRIPrep v1.3.2. Preprocessing involved generating reference volumes and skull-stripped versions using custom fMRIPrep methodology, correcting susceptibility distortions with a deformation field based on two opposing phase-encoded EPI references, and co-registering the BOLD reference to the T1w reference using nine degrees of freedom. Head-motion parameters were estimated before spatiotemporal filtering, and BOLD runs were slice-time corrected before being resampled to MNI152NLin2009cAsym standard space, resulting in preprocessed BOLD runs in this space. For analyses of whole-brain cognitive task processing without region-of-interest (ROI) analysis, images underwent additional spatial smoothing (FWHM = 4.8 mm). In all other analyses, the data was unsmoothed to prevent inclusion of extra-claustral signal in claustrum ROI voxels. Confounding time-series such as framewise displacement (FD), DVARS, and three region-wise global signals (CSF, WM, whole-brain masks) were calculated based on unsmoothed preprocessed BOLD data, along with physiological regressors for component-based noise correction.

### Cognitive Tasks

The DMCC55B protocol included four cognitive control tasks, each following a mixed block/event-related format. Trials were grouped into three long blocks lasting approximately 3 minutes each, and every block began and ended with a 30 second fixation period. The number of trials in each block was a fixed number that varied between tasks and the order of trials was randomized. There were two runs of each task, approximately 12 minutes each (24 minutes total per task). The four tasks are as described below by Etzel et al. (2022), minimally edited for style.

#### Stroop

Participants are asked to respond verbally to the ink colour of the stimulus, rather than the printed word (Stroop, 1935). The trials are termed Congruent or Incongruent based on if the printed words indicate its font colour (see Figure A1). A total of eight colours were used and divided into two sets: “MC” (mostly congruent), or “PC50” (50% Congruent). Incongruent trials were considered to have high cognitive control demands due to the conflict between colour and word dimensions, while Congruent trials were considered to have low cognitive control demands.

#### AX-CPT (Continuous Performance Task)

Participants are asked to respond to two event trials, each beginning with a Cue (a letter), followed by a Probe (a letter or number; MacDonald, 2008). Responses are a button press, the first always being button 1 in response to the Cue. The second response is button 2 if the probe is a target pair, button 1 for non-target, and no press for a No-go trial. The AX trial type (Cue A Probe X) is the target and the only condition in which the correct response is button 2. All other trial types (AY, BY, BX) are non-target, with button 1 as the respective correct response (see Figure A2). Performance is expected to vary with trial type: reaction time (RT) slower on AY and BX trials, relative to AX and BY trials, due to response conflict with the Probe that requires the use of the Cue for context.

#### Cued task-switching

Participants are prompted with a Cue to either “Attend Letter” (respond with button 1 if the letter is a vowel; button 2 if a consonant) or “Attend Number” (respond with button 1 if the number is even; button 2 if odd; Minear & Shah, 2008). The stimuli can be further described as congruent or incongruent: congruent trials being those in which the same response should be made to the Probe, irrespective of the Cue; or incongruent trials in which the correct response to the Probe depends on the Cue. Trials could be further described as repeat (when the current trial Cue is the same as that of the previous trial) or switch (when the current trial Cue is different than that of the previous trial). Incongruent trials are expected to have higher cognitive control demands due to the response conflict occurring with the Probe, and a similar pattern is expected with switch relative to repeat trials (see Figure A3).

#### Sternberg Working Memory

This version of the Sternberg working memory task uses lists with 5-8 words, split between two List screens and followed by a Probe word (Sternberg, 1966). Participants are asked to press button 2 if the Probe was in either List of the current trial, and button 1 if it was not. The Probe word was a member of the List in half of the trials (45 total), with this trial type termed NP (“novel positive”). In most of the remaining trials (36 total) the Probe is new, not in any previous List (type NN, “novel negative”). However, in 9 trials the Probe was in the immediately preceding trial’s List (type RN, “recent negative”). The trial type frequencies varied according to list length, with five-word lists having the highest frequency (see Figure A4). RN trials were considered to have high cognitive control demands, as the recent familiarity of the Probe would cause interference with processing based on working memory contents, whereas NN trials were considered to have low control demands since the Probes had no familiarity. RT was also expected to increase with List length. For the purposes of this study, List 1 and List 2 were combined into one timeframe of “Encoding”, the Delay classified as “Retention” and the Probe as “Retrieval”.

### fMRI Analysis and Statistics

To determine whole-brain task activation, first level analysis was performed for each participant to model the hemodynamic response associated with different experimental conditions using a general linear model (GLM). All models included the confounding time-series and physiological regressors mentioned above. Regressors were convolved with a canonical hemodynamic response function (HRF), and the time-series were high-pass filtered and modeled with AR(1). Initial models included all experimental conditions that showed behavioural differences, modelled across the full trial duration (from cue to probe/feedback, not including ITI). Additional models were then created that included all experimental conditions broken down into the phases of each trial.

For the Stroop task, this included 4 condition-related regressors: PC50 Congruent, PC50 Incongruent, MC Congruent and MC Incongruent; each lasting the full trial duration of 2 seconds. For the AX-CPT task, the model included 24 condition-related regressors: the 6 trial types (AX, AY, Ang, BX, BY, Bng), each broken down into three trial phases (cue, delay, probe). In this analysis, the onset of a border preceding the probe stimulus (0.3 s) was combined with the probe itself (0.5 s), totaling 0.8 s, and both together categorized as the "probe" phase. For the cued task-switching paradigm, two separate models were created to look at both (i) trial type (congruent, incongruent), and (ii) trial switch (repeat, switch). Each model included 8 condition-related regressors: the two types of conditions, each broken down into four trial phases (cue, delay, probe, feedback). For the Sternberg working memory task, the model included 9 condition-related regressors: the 3 trial types (NP, NN, RN), each broken down into three trial phases (encoding, retention, retrieval).

To ensure that multicollinearity was not a concern in our fMRI data analysis, Variance Inflation Factors (VIFs) were calculated for all the regressors of interest in our model using the Canlab toolbox (Wager, 2014). All VIF values were below a threshold of 5 (see Tables B20-22), indicating that multicollinearity was likely minimal. While a VIF below 5 does not completely rule out the presence of multicollinearity among regressors (Vatcheva et al., 2016), our relatively large sample size and the fact that VIFs remained well below the more general threshold of 10 (Hair et al., 2019), suggests that multicollinearity was not a significant concern.

#### Whole brain analyses

One sample t-tests with a cluster-forming threshold of *p* < .001 (uncorrected) were initially performed on all tasks using the smoothed images and contrast maps were generated for each task. Individual contrast maps were then entered into second-level group analyses to identify consistent activation patterns across participants. A task-specific cluster-wise threshold, determined using *p* < .001, was applied to each contrast map to correct for multiple comparisons. All whole-brain analyses were performed in MATLAB using SPM12 (https://www.fil.ion.ucl.ac.uk/spm/software/spm12/).

#### Small Region Confound Correction

To remove the effect of neighbouring insular cortex and putamen on claustrum signal, a task-adapted version of small region confound correction (SRCC) was used. As described by Stewart et al. (2024), this technique involved creating insular cortex and putamen ROIs by dilating each hemisphere’s claustrum ROI by 4 functional voxels (9.6 mm). Next, the overlap between the dilated claustrum ROIs and their neighboring insula and putamen ROIs at least 2 functional voxels (4.8 mm) away from the original claustrum were identified. This generated “flanking” ROIs within the insular cortex and putamen, similar in shape, but sufficiently distant from the claustrum to avoid including claustrum signal. The CONN Toolbox (RRID: SCR_009550; Nieto-Castanon, 2020) was used to generate canonical HRF-convolved condition time-series for each subject and task. A regressor was then generated for each flanking ROI by obtaining the interaction between the ROI de-meaned time-series and the de-meaned, summed HRF-convolved time-series of all modeled conditions per task. This procedure resulted in a time-series, which was also de-meaned, for each flanking ROI that covaried with the ROI physiological time-series during task conditions but lacked variation potentially induced by the conditions. This allowed us to control for the influence of neighboring insular and putamen regions on claustrum signal without indirectly removing task effects.

#### ROI analyses

Claustrum ROI analyses were performed by extracting the mean contrast estimate (i.e., activation value) across all voxels within the claustrum for each task condition, using MarsBaR software (Brett et al., 2002). This process was first performed to analyze claustrum activity during the whole duration of the trial, and then repeated with conditions broken down into their respective trial phases. Bilateral claustrum, putamen, and insular cortex ROIs were defined as in Stewart et al. (2024; https://identifiers.org/neurovault.collection:16667). All one-sample and paired t-tests were performed in MATLAB (https://www.mathworks.com/products/matlab.html), and the resulting *p*-values combined across all conditions and trial phases for each individual task. Multiple comparisons correction was then performed on the combined *p*-values for each task using Benjamini-Hochberg FDR correction. All reported values represent corrected *p*-values. Full tables of uncorrected and corrected *p*-values can be found in the appendix (see Tables B15-19).

Left claustrum (LCL) and Right claustrum (RCL) activation values from each task were also subjected to a repeated measures analysis of variance (ANOVA) using IBM SPSS Statistics software (version 29), with the task-specific conditions and trial phases as within-subject factors. Bonferroni correction was applied to all post-hoc tests to control for multiple comparisons. Level of significance was set at α = 0.05.

## Results

### Significant bilateral claustrum activation during high cognitive demand condition of the Stroop Task

Paired t-tests between incongruent and congruent trials indicated a difference in RT, with incongruent trials having significantly higher RTs (see Appendix A, Figure 5A). Whole-brain analysis examining incongruent over congruent conditions throughout the full trial duration revealed robust activation in task-positive regions of the frontoparietal network, including the middle and inferior frontal gyri, and superior parietal lobe. Significant activation was also seen in the occipital lobes, anterior insula, and supplementary motor cortex (Fig. 1A). Coordinates of peak cluster regions can be found in the appendix (see Table B11).

**Figure 1.**
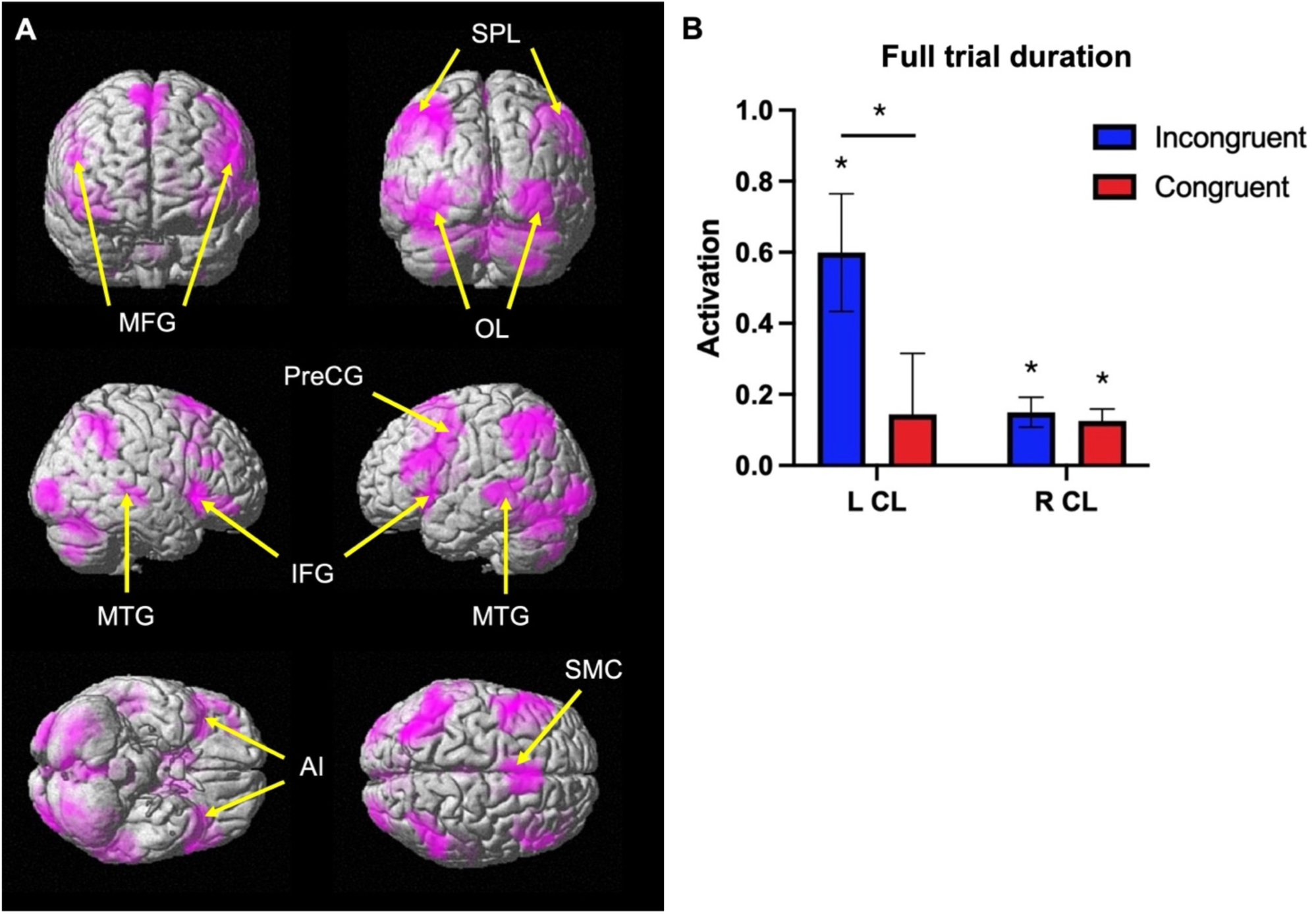
Significant task-positive and bilateral claustrum activation during high cognitive demand condition of the Stroop Task. **(A)** fMRI results displaying group-level analysis during Stroop task ([incongruent] > [congruent]). Cluster forming threshold *p* < .001 uncorrected, cluster corrected FDR (minimum cluster size = 79 voxels). SPL = superior parietal lobe; MFG = middle frontal gryus; OL = occipital lobe; PreCG = precentral gyrus; MTG = middle temporal gyrus; IFG = inferior frontal gyrus; AI = anterior insula; SMC = supplementary motor cortex. **(B)** Average LCL and RCL activation for all participants throughout whole trial duration. Error bars show standard error of the mean.

One sample t-tests across the full trial duration revealed significant bilateral claustrum activation during the incongruent condition (LCL: t = 3.62, p < .001; RCL: t = 3.58, p < .001) and significant activation during the congruent condition in the RCL (t = 3.8, p < .001), but not the LCL (t = 0.83, p = .41). Paired t-tests between incongruent and congruent conditions in the LCL showed significant differences between conditions (t = 3.72, p < .001), but not between incongruent and congruent conditions in the RCL (t = 0.88, p = .38; Fig. 1B). The results indicate significantly higher claustrum activation in the left claustrum during incongruent trials, coupled with task-positive network activation.

### Significant bilateral activation of the claustrum during cue and probe phases of the AX-CPT task

Paired t-tests between AX, BX, AY, and BY conditions indicated a difference in RT between all conditions, with RT for BX conditions being significantly higher, followed by AY conditions (see Appendix A, Figure 5B). Whole-brain analysis examining overall task effects throughout the full trial duration revealed significant task-positive activation in regions of the frontoparietal network, including the middle and inferior frontal gyri, and superior parietal lobe. Significant activations can also be seen in regions such as the occipital lobes, pre- and post-central gyrus, anterior insula, and supplementary motor cortex (Fig. 2A). Coordinates of peak cluster regions can be found in the appendix (see Table B12).

**Figure 2.**
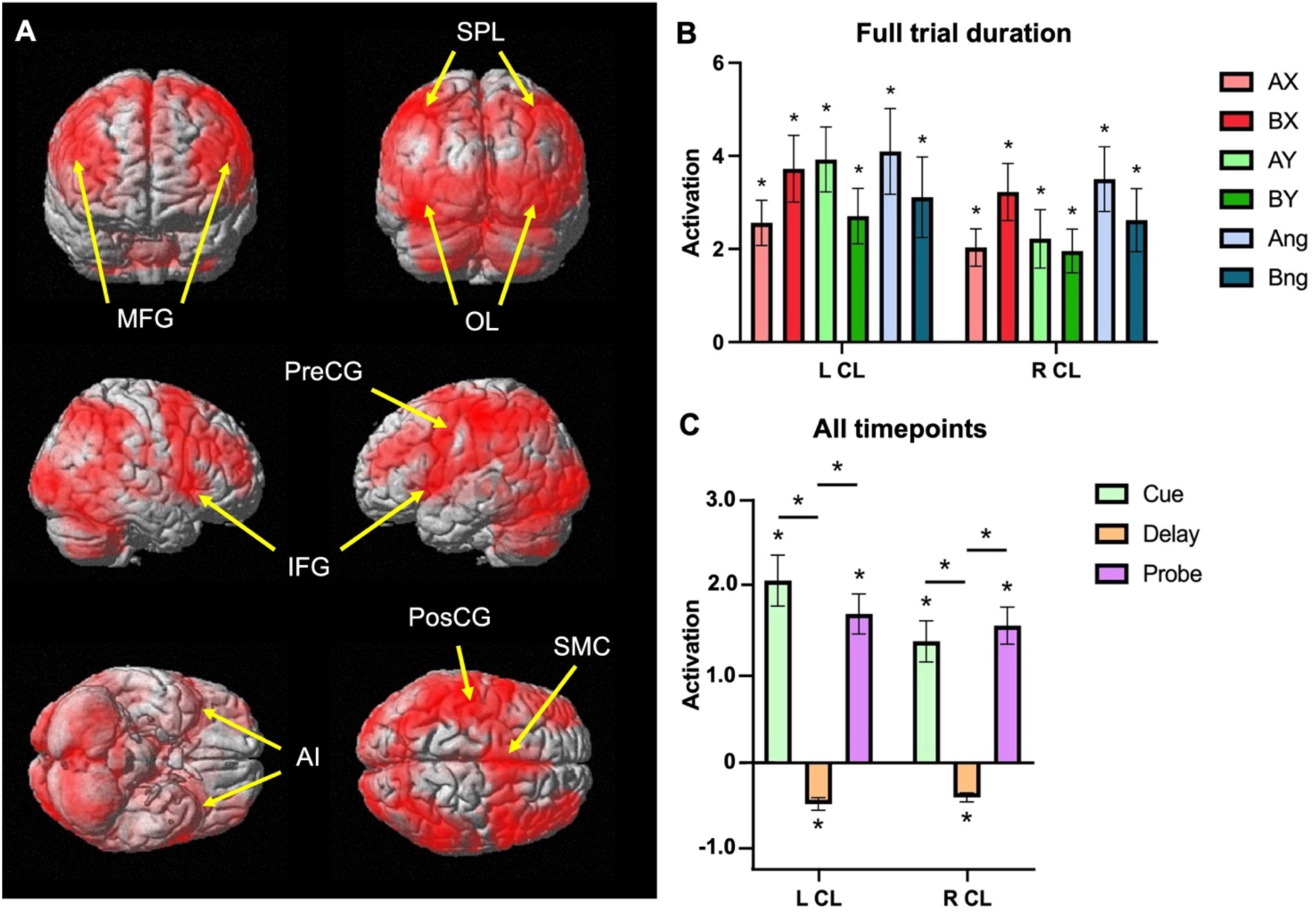
Significant task-positive and bilateral activation of the claustrum during cue and probe phases of the AX-CPT task. **(A)** fMRI results displaying group-level analysis during AX-CPT task ([AX, AY, Ang, BX, BY, Bng] > [baseline]). Cluster forming threshold *p* < .001 uncorrected, cluster corrected FDR (minimum cluster size = 49,272 voxels). SPL = superior parietal lobe; MFG = middle frontal gryus; OL = occipital lobe; IFG = inferior frontal gyrus; PreCG = precentral gyrus; PosCG = postcentral gyrus; AI = anterior insula; SMC = supplementary motor cortex. **(B)** Average LCL and RCL claustrum activation for all participants throughout whole trial duration **(C)** Average LCL and RCL activation during trial phases, pooling all conditions. Error bars show standard error of the mean.

One sample t-tests across the full trial duration revealed significant activation of both the LCL (AX: t = 5.30, p < .001; BX: t = 5.20, p < .001; AY: t = 5.61, p < .001; BY: t = 4.56, p < .001; Ang: t = 4.45, p < .001; Bng: t = 3.60, p < .001) and RCL (AX: t = 5.09, p < .001; BX: t = 5.25, p < .001; AY: t = 3.55, p < .001; BY: t = 4.18, p < .001; Ang: t = 5.05, p < .001; Bng: t = 3.87, p < .001) during all trial types (Fig. 2B). When examining claustrum activation by trial phases during the “Cue” (0.5 s), “Delay” (4 s) and “Probe” (0.8 s), one-sample t-tests revealed significant bilateral claustrum activation during the cue (LCL: t = 7.16, p < .001; RCL: t = 5.94, p < .001) and probe (LCL: t = 7.44, p < .001; RCL: t = 7.44, p < .001) phases, as well as significant bilateral deactivation during the delay (LCL: t = -6.51, p < .001; RCL: t = -7.69, p < .001; Fig. 2C).

Examining each phase separately by trial type, one-sample t-tests revealed significant activation of both the LCL (AX: t = 4.94, p < .001; BX: t = 4.70, p < .001; AY: t = 4.65, p < .001; BY: t = 3.72, p < .001; Ang: t = 4.19, p < .001; Bng: t = 3.52, p < .001) and RCL (AX: t = 4.13, p < .001; BX: t = 4.22, p < .001; AY: t = 2.12, p = .038; BY: t = 3.45, p = .001; Ang: t = 3.93, p < .001; Bng: t = 3.04, p = .004) for all trial types during the cue. Significant activation of both LCL (AX: t = 5.99, p < .001; BX: t = 4.33, p < .001; AY: t = 4.83, p < .001; BY: t = 4.82, p < .001; Ang: t = 5.78, p < .001; Bng: t = 4.56, p < .001) and RCL (AX: t = 6.15, p < .001; BX: t = 5.52, p < .001; AY: t = 4.68, p < .001; BY: t = 5.24, p < .001; Ang: t = 5.70, p < .001; Bng: t = 4.69, p < .001) for all trial types was also seen during the probe. Significant deactivation of both LCL (AX: t = -3.52, p < .001; BX: t = -3.64, p < .001; AY: t = -4.33, p < .001; BY: t = -6.03, p < .001; Ang: t = -4.56, p < .001; Bng: t = -3.87, p < .001) and RCL (AX: t = -5.14, p < .001; BX: t = -5.31, p < .001; AY: t = -3.08, p < .001; BY: t = -4.94, p < .001; Ang: t = -5.67, p < .001; Bng: t = -3.98, p < .001) for all trial types was seen during the delay.

A repeated measures 6 × 3 ANOVA with two within-subject factors: condition (AX, BX, AY, BY, Ang, Bng) and trial phase (cue, delay, probe), was conducted for LCL and RCL activation to examine main effects between conditions and phases. For the LCL, a significant main effect of condition was observed, *F*_(4.35, 234.75)_ = 2.42, *p* = .044, *η^2^* = 0.043. However, post-hoc comparisons indicated that none of the pairwise comparisons between conditions were statistically significant (*p* > .05 for all comparisons). A significant main effect of trial phase was also found in the LCL, *F*_(1.81, 97.90)_ = 43.12, *p* < .001, *η^2^* = 0.444. Post-hoc comparisons between the three trial phases revealed significant activation differences between the delay and other two phases: cue and delay: *M_difference_* = 2.49, *p* < .001; delay and probe: *M_difference_* = 2.02, *p* < .001. Although there was no significant interaction effect between condition and trial phase, *F*_(6.22, 335.73)_ = 1.42, *p* = .21, *η^2^* = 0.026, differences across trial phases within each condition likely contributed to the main effect of condition.

For the RCL, no main effect of condition was observed, *F*_(4.37, 235.95)_ = 1.76, *p* = .13, *η^2^* = 0.031. However, a significant main effect of trial phase was observed, *F*_(3, 324)_ = 44.55, *p* < .001, *η^2^* = 0.452. Post-hoc comparisons between the three phases revealed significant activation differences similar to those observed in the LCL: cue and delay: *M_difference_* = 1.80, *p* < .001; delay and probe: *M_difference_* = 1.98, *p* < .001. Overall, these findings highlight the significant activation of the claustrum during all trial conditions, particularly during the cue and probe phases, coupled with task-positive network activation.

### Significant bilateral claustrum activation during the delay phase of cued task-switching

Paired t-tests between incongruent and congruent trials indicated a difference in RT, with incongruent trials having significantly higher RTs. Likewise for switch compared to repeat trials, paired t-tests revealed differences in RT, with switch trials having higher RTs (see Appendix A, Figure 5C). Whole-brain analysis examining overall task effects throughout the full trial duration revealed activation in areas such as the occipital lobes, superior and middle temporal gyri, precentral gyrus, and supplementary motor cortex (Fig. 3A). Coordinates of peak cluster regions can be found in the appendix (see Table B13).

**Figure 3.**
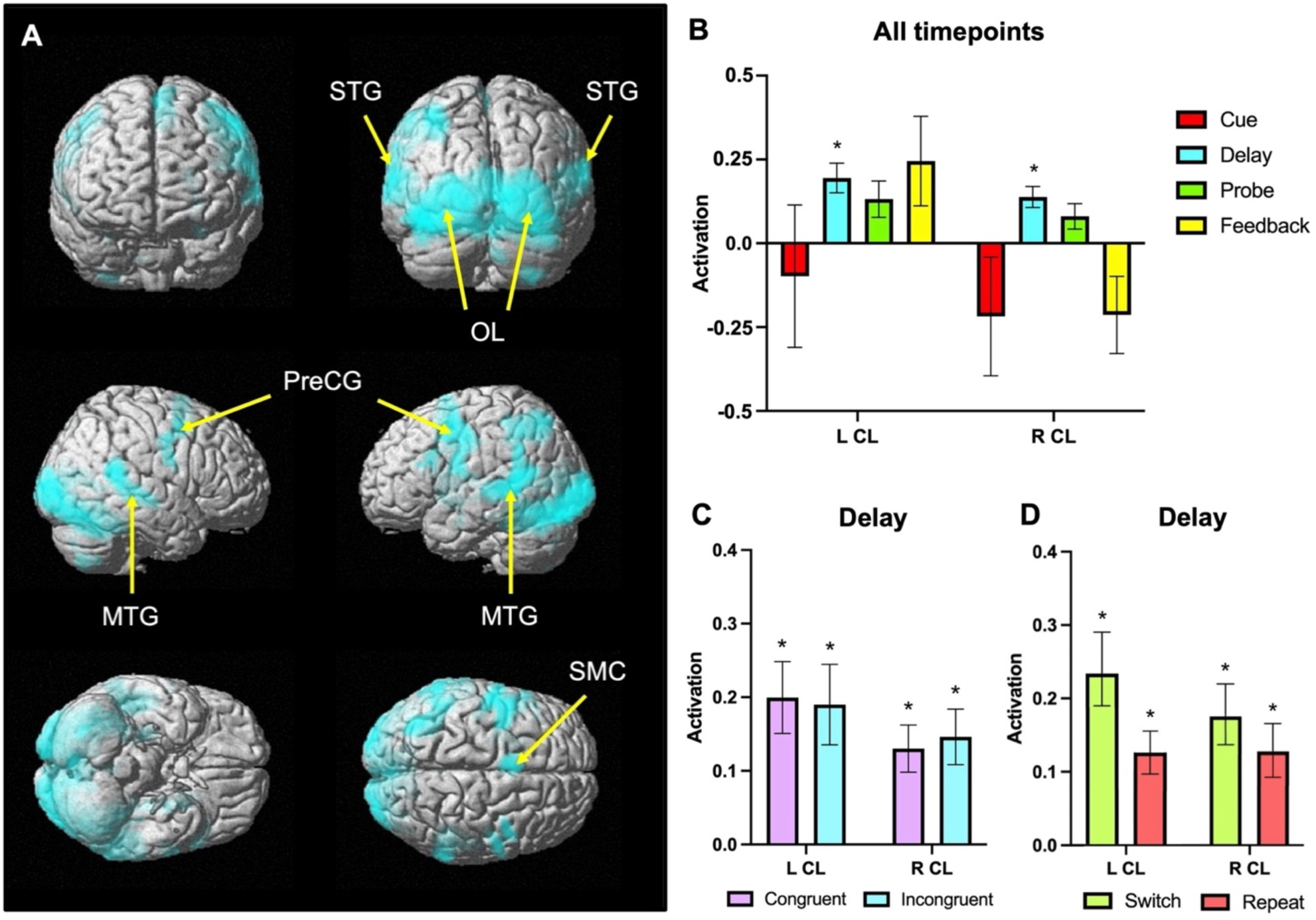
Significant bilateral claustrum activation during delay phase of cued task-switching. **(A)** fMRI results displaying group-level analysis during cued task-switching ([congruent, incongruent] > [baseline]). Cluster forming threshold *p* < .001 uncorrected, cluster corrected FDR (minimum cluster size = 66 voxels). STG = superior temporal gyrus; OL occipital lobe; MTG = middle temporal gyrus; PreCG = precentral gyrus; SMC = supplementary motor cortex. **(B)** Average LCL and RCL activation during trial phases, pooling all conditions. **(C)** Average LCL and RCL activation by trial type during delay phase. **(D)** Average LCL and RCL activation by trial switch during delay phase. Error bars show standard error of the mean.

One sample t-tests revealed no significant activation for either condition in the LCL (Congruent: t = 1.22, p = .32; Incongruent: t = 1.99, p = .12) or RCL (Congruent: t = - 1.02, p = .38; Incongruent: t = -1.06, p = .37) when broken down by trial type. Similarly, no significant activation was found for either condition in the LCL (Switch: t = 1.45, p = .26; Repeat: t = 1.26, p = .33) or RCL (Switch: t = -1.50, p = .26; Repeat: t = -0.90, p = .46) when broken down by trial switch. No significant difference was found between the conditions in either model. When examining claustrum activation by trial phases during the “Cue” (0.5 s), “Delay” (4 s), “Probe” (3.2 s) and “Feedback” (0.8 s), one-sample t-tests revealed significant bilateral claustrum activation only during the delay portions of each trial (LCL: t = 4.41, p < .001; RCL: t = 4.44, p < .001; Fig. 3B).

Examining each phase separately by trial type, one sample t-tests revealed significant bilateral activation for both conditions (Congruent LCL: t = 4.10, p = .001; Incongruent LCL: t = 3.48, p = .005; Congruent RCL: t = 4.08, p = .001; Incongruent RCL: t = 3.87, p = .002) during the delay (Fig. 3C). The trial switch model followed the same trends of activation as the trial type model, with significant bilateral activation seen for both switch and repeat trials during the delay (Switch LCL: t = 4.33, p < .001; Repeat LCL: t = 3.49, p = .005; Switch RCL: t = 4.22, p < .001; Repeat RCL: t = 3.68, p = .004; Fig. 3D).

A repeated measures 2 × 4 ANOVA with two within-subject factors: condition (congruent, incongruent) and trial phase (cue, delay, probe, feedback), was conducted for LCL and RCL activation to examine main effects between trial type and phases. For the LCL, no significant main effect of condition, *F*_(1, 54)_ = 1.58, *p* = .21, *η^2^* = 0.033, nor of trial phase, *F*_(1.46, 78.83)_ = 1.36, *p* = .26, *η^2^* = 0.025, was found. Similarly, for the RCL, no significant main effect of condition, *F*_(1, 54)_ = .04, *p* = .83, *η^2^* = 0.001, nor of trial phase, *F*_(1.50, 81.07)_ = 2.81, *p* = .081, *η^2^* = 0.049, was found.

A separate repeated measures 2 × 4 ANOVA with two within-subject factors: condition (switch, repeat) and trial phase (cue, delay, probe, feedback), was also conducted to examine main effects between trial switch and phases. For the LCL, no significant main effect of condition, *F*_(1, 54)_ = 0.27, *p* = .61, *η^2^* = 0.005, nor of trial phase, *F*_(1.46, 78.76)_ = 1.51, *p* = .23, *η^2^* = 0.027, was found. Similarly, for the RCL, no significant main effect of condition, *F*_(1, 54)_ = 0.54, *p* = .47, *η^2^* = 0.01, nor of trial phase, *F*_(1.45, 78.21)_ = 3.00, *p* = .072, *η^2^* = 0.053, was found. These results indicate significant bilateral claustrum activation during the delay phase across both trial type and trial switch, with no significant claustrum activation differences between conditions.

### Significant bilateral claustrum activation during retrieval phase of the Sternberg working memory task

Paired t-tests between all trial types revealed significant differences in RT, with RN trials having significantly higher RTs than NN and NP trials (see Appendix A, Figure 5D). Paired t-tests between list lengths revealed no significant differences in RT (see Appendix A, Figure 5E), thus differences were only looked at between trial types.

Whole-brain analysis examining overall task effects throughout the full trial duration revealed significant task-positive activation in regions of the frontoparietal network, including the middle and inferior frontal gyri, and superior parietal lobe (Fig. 4A). Significant activations could also be seen in regions such as the occipital lobes, precentral gyrus, anterior insula, and supplementary motor cortex. Coordinates of peak cluster regions can be found in the appendix (see Table B14).

**Figure 4.**
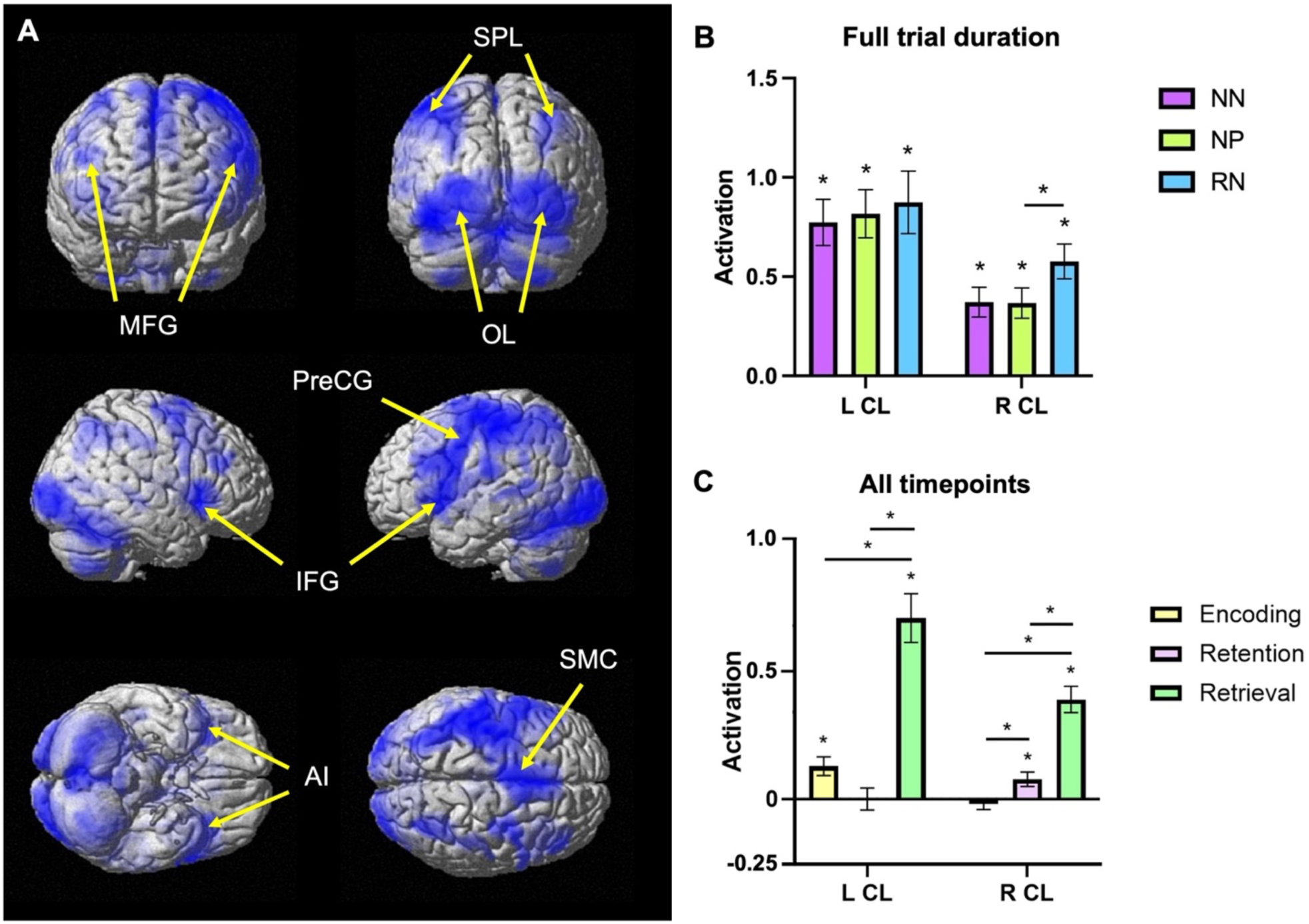
Significant task-positive and bilateral claustrum activation during retrieval phase of the Sternberg working memory task. **(A)** fMRI results displaying group-level analysis during Sternberg working memory task ([RN, NN, NP] > [baseline]). Cluster forming threshold *p* < .001 uncorrected, cluster corrected FDR (minimum cluster size = 89 voxels). SPL = superior parietal lobe; MFG = middle frontal gyrus; OL = occipital lobe; PreCG = precentral gyrus; IFG = inferior frontal gyrus; AI = anterior insula; SMC = supplementary motor cortex. **(B)** Average LCL and RCL activation for all participants throughout whole trial duration. **(C)** Average LCL and RCL activation during trial phases, pooling all conditions. Error bars show standard error of the mean.

One sample t-tests revealed significant bilateral claustrum activation during all trial types when averaged over the full duration of the trial (NN LCL: t = 6.64, p < .001; NP LCL: t = 6.72, p < .001; RN LCL: t = 5.54, p < .001; NN RCL: t = 4.95, p < .001; NP RCL: t = 4.82, p < .001, RN RCL: t = 6.58, p < .001; Fig. 4B). When examining claustrum activation by trial phases during “Encoding” (4.8 s), “Retention” (4 s) and “Retrieval” (1.6 s), one sample t-tests revealed significant activation in the LCL during the encoding phase (t = 3.49, p = .002), in the RCL during the retention phase (t = 2.70, p = .015), and in bilateral claustrum during the retrieval phase (LCL: t = 7.42, p < .001; RCL: t = 7.53, p < .001; Fig. 4C).

Examining each phase separately by trial type, one-sample t-tests revealed that significant activation in the LCL during the encoding phase was attributed to the NP and RN conditions (NP: t = 2.86, p = .010; RN: t = 3.87, p < .001), with no significant activation in the RCL. Significant activation seen in the RCL during the retention phase was attributed to the NN and NP trials (NN: t = 2.39, p = .031; NP: t = 3.83, p < .001), with no significant activation seen in the LCL. Bilateral claustrum showed significant activation of all trial types during the retrieval phase (NN LCL: t = 6.80, p < .001; NP LCL: t = 7.13, p < .001; RN LCL: t = 4.42, p < .001; NN RCL: t = 5.31, p < .001; NP RCL: t = 4.72, p < .001; RN RCL: t = 5.83, p < .001).

A repeated measures 3 × 3 ANOVA with two within-subject factors: condition (NN, NP, RN) and trial phase (encoding, retention, retrieval), was conducted for LCL and RCL activation to examine effects between conditions and phases. For the LCL, no significant main effect of condition was observed, *F*_(1.63, 87.88)_ = 0.38, *p* = .64, *η^2^* = 0.007. However, a significant main effect of trial phase was found, *F*_(1.28, 68.83)_ = 36.97, *p* < .001, *η^2^* = 0.406. Post-hoc comparisons between the three phases revealed significant activation differences between retrieval and the other two phases: encoding and retrieval: *M_difference_* = 0.57, *p* < .001; retention and retrieval: *M_difference_* = 0.70, *p* < .001 (Fig. 4C).

For the RCL, a main effect of condition was observed, *F*_(1.48, 79.73)_ = 5.03, *p* = .016, *η^2^* = 0.085, with post-hoc comparisons indicating significant activation differences between NP and RN trials: *M_difference_* = 0.07, *p* = .036 (Fig. 4B). A significant main effect of trial phase was also observed, *F*_(1.51, 81.49)_ = 37.97, *p* < .001, *η^2^* = 0.413, with post-hoc comparisons revealing significant activation differences between all phases: encoding and retention: *M_difference_* = 0.10, *p* = .028; encoding and retrieval: *M_difference_* = 0.40, *p* < .001; retention and retrieval: *M_difference_* = 0.31, *p* < .001 (Fig. 4C). Additionally, a significant interaction effect between condition and trial phase was found, *F*_(1.73, 93.41)_ = 3.43, *p* = .043, *η^2^* = 0.060. The results indicate significant bilateral claustrum activation observed across all trial types, particularly during the retrieval phase, alongside task-positive network activation.

## Discussion

In this study, we analyzed human fMRI data involving four different cognitive tasks to investigate the relationship between the claustrum and cognitive control. We isolated a distinct claustrum signal by using a task-adapted version of SRCC that minimized the removal of task effects (Stewart et al., 2024), and examined claustrum activation during distinct phases of each trial. Whole brain analyses of the four tasks displayed activity in the frontoparietal network and other sensory and association cortices, confirming the activation of task-positive networks. Claustrum activity was present during all four tasks, with activation peaking at certain points of each trial duration. The results indicate that claustrum activity is associated with tasks that activate task-positive networks, particularly during times of increased cognitive demand and active use of cognitive control.

The Stroop task showed activity consistent with our hypothesis, revealing notable claustrum activation during incongruent trials—particularly in the left hemisphere—as compared to congruent trials. The behavioural results indicated heightened cognitive demand during incongruent trials, and this was paralleled by markedly increased claustrum activation. The AX-CPT and Sternberg tasks both revealed significant bilateral claustrum activation during all conditions when examining the full trial duration, as well as a significant main effect of trial phase during both tasks. Heightened claustrum activation was seen particularly during the cue and probe phases in the AX-CPT task, coupled with deactivation during the delay phase. The highest activation for all conditions in the Sternberg task was seen during the retrieval phase.

Evidence from these three tasks indicate heightened claustrum activation during trial phases requiring immediate and meaningful responses, suggesting that the claustrum becomes particularly engaged when increased cognitive control is necessary to process and respond to meaningful stimuli. All three tasks also exhibited strong task-positive network activation in characteristic regions associated with task-related activations, such as frontoparietal areas, bilateral anterior insula, and supplementary motor cortex (Biswal et al., 1995; Broyd et al., 2009; Fox et al., 2005). These results are consistent with the theory that the claustrum functions as a network instantiator, indicating that heightened activation during each task reflects a period of increased cognitive demand during which the claustrum becomes engaged (Madden et al., 2022).

Despite the overall high claustrum activation during each condition, there was no difference between any of the trial types in the AX-CPT task. Behaviourally, the AY and BX conditions showed greater cognitive demand compared to the AX and BY conditions, suggesting that we would observe heightened claustrum activity during these conditions. However, the prediction that certain conditions would show stronger activation was based on limited prior evidence from the multi-source interference task (MSIT; Krimmel et al., 2019; Stewart et al., 2024). Given that claustrum activation has not been extensively studied in tasks beyond the MSIT, the widespread activation observed in all conditions may reflect the generally high cognitive demand associated with the AX-CPT task. Similarly, during the Sternberg task, higher claustrum activity was predicted during RN trials, compared to NN or NP. This pattern was observed in the RCL, however, the increased activation during RN trials should be interpreted with caution, as this finding may be affected by limited statistical power due to the smaller number of RN trials (9), compared to NN (36) and NP (45) trials.

During the cued task-switching paradigm, the claustrum exhibited activation solely during the delay phase. Unlike the previous tasks where the cue requires immediate action or memory retention, this cue primarily prepares the participant for the upcoming stimulus and provides information on where to direct their attention. This could potentially explain why the claustrum displays heightened engagement during the delay phase; it may be actively involved in preparing and aligning cognitive resources in anticipation of the upcoming stimulus. Notably, the smaller magnitude of RT differences in this task, compared to those observed in the Stroop or AX-CPT (see Tables B2, B4, B10), coupled with the distinct lack of prefrontal cortex activation, suggests that this particular task may not be optimal for investigating claustrum activity as cognitive demand between conditions in this task is relatively low.

Throughout many of the tasks, we also observed differences in laterality of claustrum activation. Although incongruent trials in the Stroop task exhibited significantly higher activation than congruent trials in the LCL, the RCL did not follow this pattern. The tendency for greater activation in the left claustrum during cognitive tasks aligns with prior research suggesting laterality in claustrum function (Barrett et al., 2020; Krimmel et al., 2019; Stewart et al., 2024), although the reason for this functional difference remains unclear. Significant activation was also seen in the LCL during encoding of the Sternberg task, though not the RCL. This lateralization could be linked to the linguistic nature of the task, as strong activation was also seen in the left angular and supramarginal gyri, regions known to be involved in language (Price, 2012), semantics (Binder et al., 2009; Binder & Desai, 2011; Noonan et al., 2013), and phonological processing (Booth et al., 2002; Price et al., 1997). Given the extensive connections of the claustrum (Torgerson et al., 2015), the LCL’s involvement could be in helping maintain the network integrity required for semantic working memory. Interestingly, similar activations were seen during the Stroop task, which also involved whole-word stimuli. However, the Stroop, unlike the Sternberg task, required an immediate response and did not involve distinct working memory phases, preventing us from observing whether this lateralization followed the same patterns of trial phase activation as the Sternberg.

Significant activation was also seen in the RCL during the retention phase of the Sternberg task. Historically, the right hemisphere has typically been implicated in spatial working memory (Smith et al., 1996; Smith & Jonides, 1998), but recent studies have demonstrated the importance of right hemisphere involvement in verbal working memory domains as well (Ferber et al., 2020; Marti et al., 2024). Marti et al. (2024) demonstrated that optimal working memory performance, particularly the updating of verbal information, relies on the right hemisphere, notably the right frontal cortex. The RCL’s heightened engagement may therefore reflect a more dominant role in facilitating and initiating these processes, particularly during the retention phase.

## Limitations

The emergence of strong task-positive cortical networks during tasks with high claustrum activation supports the link between claustrum activity and cognitive processes; however, the potential influence of motor contributions cannot be entirely ruled out. Additional evidence from White et al. (2020) demonstrates that in mice, the claustrum is crucial for optimal performance on the 5-choice serial response time task, a task that retains the motor demands of the simpler 1-choice task, while requiring more cognitive demand. Previous functional studies investigating claustrum activity have also used the MSIT, which includes a control “tapping” task that involves the same motor movements, but lacks the cognitive processing required by the actual task (Krimmel et al., 2019; Stewart et al., 2024). Despite this, there is still evidence that the claustrum is implicated in higher-order premotor circuits, playing a role in encoding movement direction and modulating cortical activity during perceptual decision-making (Chevée et al., 2022). The implementation of motor control conditions in future cognitive tasks investigating the claustrum is crucial to eliminating confounding motor effects.

The absence of functional connectivity imaging in this study also limits the ability to assess the directional influence of the claustrum on specific networks or make claims on causal relationships. Despite this, the primary aim of the current study was to characterize claustrum activity in functional human data during tasks beyond the MSIT. To build on these findings, future research could employ functional connectivity analysis and dynamic causal modeling to explore the directional interactions between the claustrum and related brain networks, potentially providing a deeper understanding of the claustrum’s role in cognitive processes.

## Conclusion

In summary, our findings demonstrate that claustrum activation is linked to task-positive network engagement across a variety of cognitive tasks, with notable activation during periods of heightened cognitive demand that require active use of cognitive control. These results provide evidence supporting the NICC model of claustrum function and enhances our comprehension of the neural mechanisms underlying cognitive functioning. Claustrum involvement is evident in various neuropsychiatric conditions (Arrigo et al., 2019; Chen et al., 2023; Nikolenko et al., 2021). This study paves the way for an understanding of how claustrum pathology may contribute to the cognitive impairment component across neuropsychiatric conditions.

## Data Availability

The Dual Mechanisms of Cognitive Control (DMCC55B) dataset is publicly available at OpenNeuro (Braver et al., 2021). ROI files required for performing SRCC can be found at Neurovault (https://neurovault.org/collections/16667/).

## Author Contribution

Conceptualization: C.H. and D.A.S.; methodology: C.H., B.W.S., and D.A.S.; formal analysis, visualization, and writing – original draft: C.H.; writing – review & editing: C.H., B.W.S., C.L., P.P., B.N.M., and D.A.S.; supervision, project administration, and funding acquisition: D.A.S.

## Acknowledgements

The authors have no conflicts of interest to declare. Funding was provided through a Natural Sciences and Engineering Research Council of Canada Discovery Grant to DAS. Other support provided by the Wolfe-Western Fellowship At-Large for Outstanding Newly Recruited Research Scholars Endowment Fund to DAS.

