## Appendices A and B for "The human claustrum activates across multiple cognitive tasks"

### Supplemental Materials

### APPENDIX A

**Figure 1.**

*Stroop task timing, trial types and correct responses.*

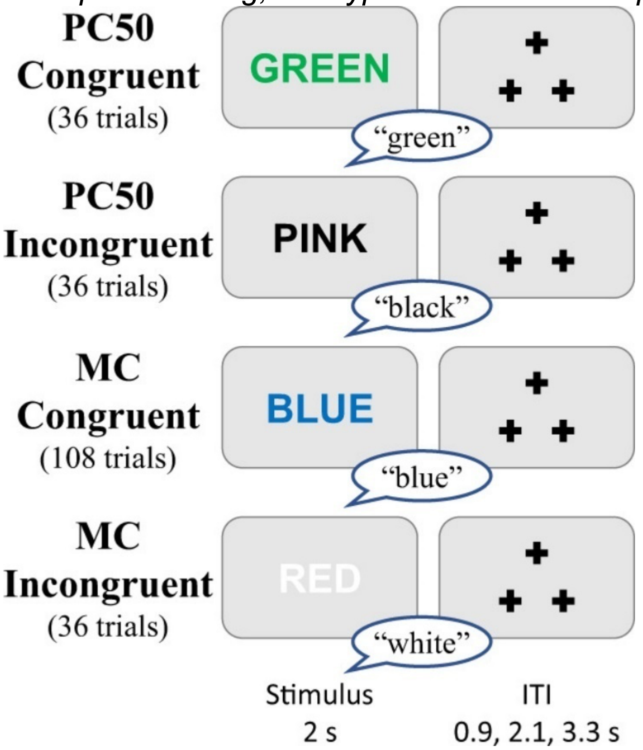


*Note*. Total number of trials of each type completed by each participant listed underneath the trial type. Figure reprinted from Etzel et al., (2022), licensed under Creative Commons Attribution 4.0 International License with permission from the authors.

**Figure 2.**

*AX-CPT task timing, trial types and correct responses.*

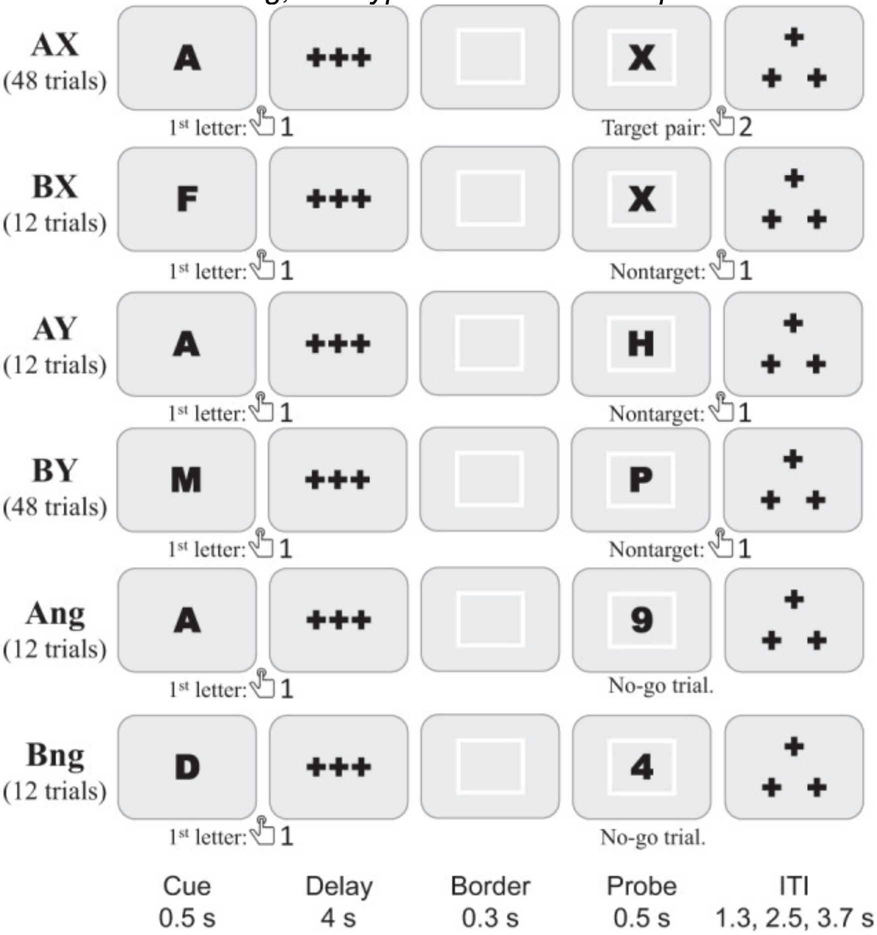


*Note*. Total number of trials of each type completed by each participant listed underneath the trial type. Figure reprinted from Etzel et al., (2022), licensed under Creative Commons Attribution 4.0 International License with permission from the authors.

**Figure 3.**

*Cued task-switching task timing, trial types and correct responses.*

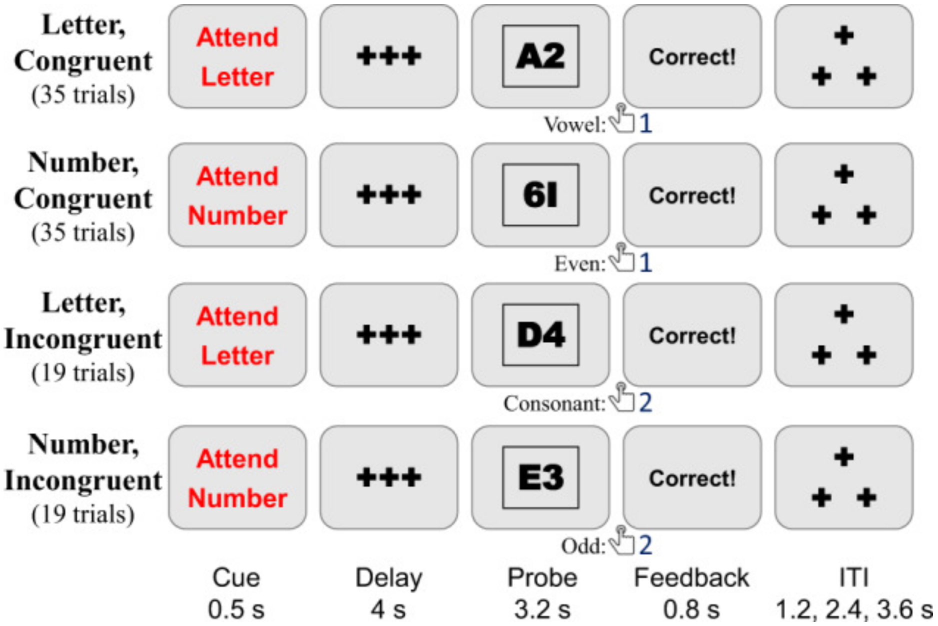


*Note*. Total number of trials of each type completed by each participant listed underneath the trial type. Figure reprinted from Etzel et al., (2022), licensed under Creative Commons Attribution 4.0 International License with permission from the authors.

**Figure 4.**

*Sternberg task timing, trial types and correct responses.*

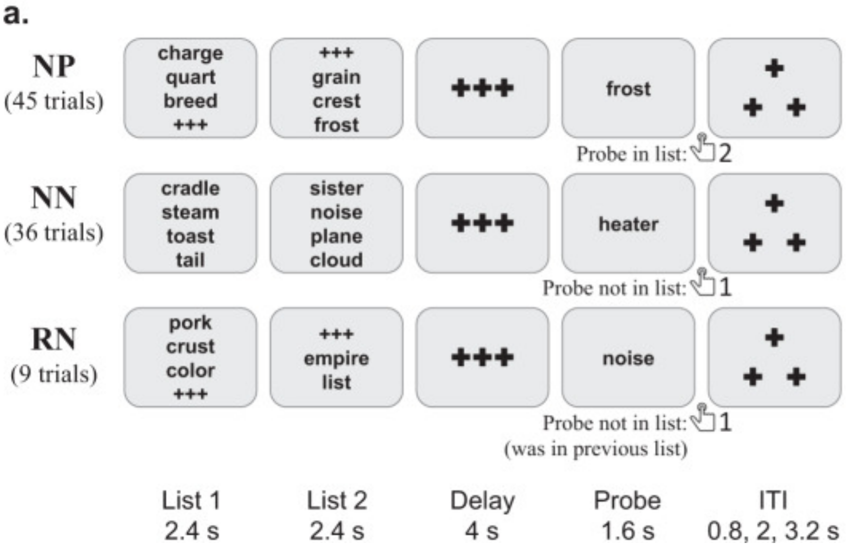


b.

| LL | NP | NN | RN |
|---|---|---|---|
| 5 | 18 | 12 | 6 |
| 6 | 4 | 5 | 0 |
| 7 | 9 | 8 | 1 |
| 8 | 14 | 11 | 2 |
| *total* | *45* | *36* | *9* |

*Note.* (a) Total number of trials of each type completed by each participant listed underneath the trial type. (b) Total number of examples of each type and list length of both lists combined. Figure reprinted from Etzel et al., (2022), licensed under Creative Commons Attribution 4.0 International License with permission from the authors.

**Figure 5.**

*Mean RT for all tasks.*

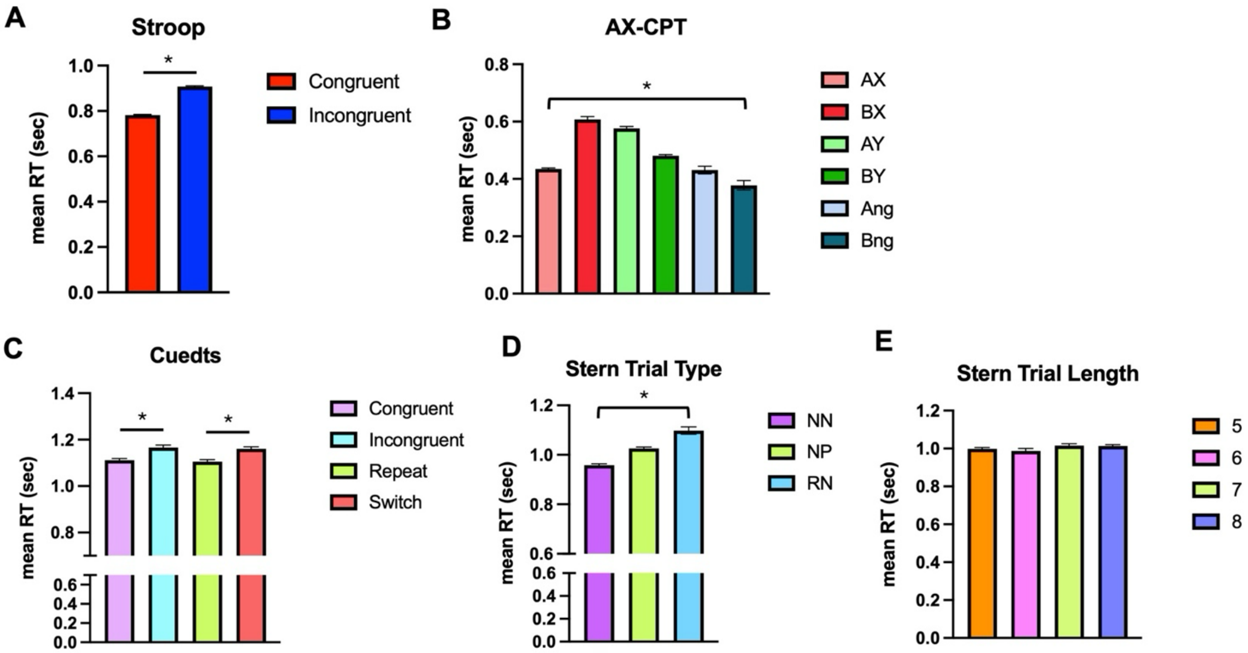


*Note.* **(A)** Stroop, **(B)** AX-CPT, **(C)** cued task-switching, **(D)** Sternberg (by trial type) and **(E)** Sternberg (by trial length). Reaction times calculated from correct and incorrect trials. Error bars show standard error of the mean.

### APPENDIX B

**Table 1.**
*Mean reaction time and error rate across trial types of Stroop.*

| Trial Type | Mean Reaction Time | RT.SEM | Mean Error Rate | ER.SEM |
|---|---|---|---|---|
| Con | 0.782 | 0.002 | 0.003 | 0.001 |
| InCon | 0.908 | 0.003 | 0.026 | 0.003 |

**Table 2.**
*Paired t-tests for RT between all combinations of trial types across all trials and participants of Stroop.*

| Comparison | P-value | T-statistic | DF |
|---|---|---|---|
| Con vs InCon | 3E-227 | -32.907 | 11757 |

**Table 3.**
*Mean reaction time and error rate across trial types of AX-CPT.*

| Trial Type | Mean Reaction Time | RT.SEM | Mean Error rate | ER.SEM |
|---|---|---|---|---|
| AX | 0.437 | 0.014 | 0.048 | 0.004 |
| AY | 0.579 | 0.015 | 0.048 | 0.008 |
| Ang | 0.437 | 0.021 | 0.111 | 0.012 |
| BX | 0.612 | 0.022 | 0.162 | 0.014 |
| BY | 0.484 | 0.016 | 0.028 | 0.003 |
| Bng | 0.396 | 0.020 | 0.155 | 0.014 |

*Note*. Ang and Bng reaction times only from incorrect trials; for all other trial types, reaction times were calculated from both incorrect and correct trials.

**Table 4.**
*Paired t-tests for RT between all combinations of trial types across all trials and participants of AX-CPT.*

| Comparison | P-value | T-statistic | DF |
|---|---|---|---|
| AX vs AY | 1.231E-80 | -19.559 | 3259 |
| AX vs Ang | 0.829 | 0.216 | 2678 |
| AX vs BX | 2.627E-97 | -21.662 | 3249 |

| | | | |
|---|---|---|---|
| AX vs BY | 1.524E-22 | -9.815 | 5203 |
| AX vs Bng | 4.268E-04 | 3.527 | 2707 |
| AY vs Ang | 5.003E-11 | 6.672 | 725 |
| AY vs BX | 8.996E-03 | -2.616 | 1296 |
| AY vs BY | 6.583E-33 | 12.082 | 3250 |
| AY vs Bng | 1.219E-23 | 10.369 | 754 |
| Ang vs BX | 2.814E-09 | -6.018 | 715 |
| Ang vs BY | 0.170 | -2.389 | 2669 |
| Ang vs Bng | 0.187 | 2.373 | 173 |
| BX vs BY | 8.780E-48 | 14.763 | 3240 |
| BX vs Bng | 1.189E-18 | 9.054 | 744 |
| BY vs Bng | 1.033E-08 | 5.743 | 2698 |

**Table 5.**

*Mean reaction time and error rate across trial types of Sternberg.*

| Trial Type | Mean Reaction Time | RT.SEM | Mean Error Rate | ER.SEM |
|---|---|---|---|---|
| NN | 0.958 | 0.006 | 0.059 | 0.005 |
| NP | 1.026 | 0.006 | 0.147 | 0.007 |
| RN | 1.098 | 0.015 | 0.317 | 0.021 |

**Table 6.**

*Mean reaction time and error rate across trial lengths of Sternberg.*

| Trial Length | Mean Reaction Time | RT.SEM | Mean Error Rate | ER.SEM |
|---|---|---|---|---|
| 5 | 0.999 | 0.007 | 0.106 | 0.007 |
| 6 | 0.987 | 0.013 | 0.113 | 0.014 |
| 7 | 1.016 | 0.009 | 0.162 | 0.012 |
| 8 | 1.013 | 0.007 | 0.142 | 0.009 |

**Table 7.**

*Paired t-tests for RT between all combinations of trial types and lengths across all trials and participants of Sternberg.*

| Comparison | P-value | T-statistic | DF |
|---|---|---|---|
| NN vs NP | 2.890E-14 | -7.629 | 4363 |
| NN vs RN | 1.935E-21 | -9.599 | 2416 |
| NP vs RN | 3.081E-06 | -4.675 | 2887 |
| 5 vs 6 | 0.467 | 0.728 | 2420 |
| 5 vs 7 | 0.155 | -1.424 | 2897 |
| 5 vs 8 | 0.163 | -1.396 | 3381 |
| 6 vs 7 | 0.078 | -1.761 | 1451 |
| 6 vs 8 | 0.084 | -1.730 | 1935 |
| 7 vs 8 | 0.831 | 0.213 | 2412 |

**Table 8.**

*Mean reaction time and error rate across trial types of cued task-switching.*

| Trial Type | Mean Reaction Time | RT.SEM | Mean Error Rate | ER.SEM |
|---|---|---|---|---|
| Congruent | 1.112 | 0.007 | 0.033 | 0.003 |
| Incongruent | 1.166 | 0.011 | 0.075 | 0.006 |

**Table 9.**

*Mean reaction time and error rate across trial switch of cued task-switching.*

| Trial Switch | Mean Reaction Time | RT.SEM | Mean Error Rate | ER.SEM |
|---|---|---|---|---|
| n/a | 1.068 | 0.023 | 0.036 | 0.01 |
| repeat | 1.105 | 0.009 | 0.041 | 0.004 |
| switch | 1.161 | 0.009 | 0.055 | 0.004 |

**Table 10.**

*Paired t-tests for RT between all combinations of trial types and switches across all trials and participants of cued task-switching.*

| Comparison | P-value | T-statistic | DF |
|---|---|---|---|
| Congruent vs Incongruent | 1.404E-05 | -4.347 | 5908 |
| n/a vs repeat | 0.156 | -1.419 | 2969 |
| n/a vs switch | 7.068E-04 | -3.390 | 3263 |
| repeat vs switch | 9.934E-06 | -4.423 | 5582 |

**Table 11.**

*Peak clusters of brain regions exhibiting significant increases in activation during Stroop.*

| Region | x (mm) | y (mm) | z (mm) |
|---|---|---|---|
| Left Angular Gyrus | -34 | 65 | 47 |
| Right Angular Gyrus | 46 | -55 | 54 |
| Left Caudate | 14 | 2 | 16 |
| Right Caudate | 12 | 12 | 4 |
| Left Precentral Gyrus | -46 | 7 | 32 |
| Left Anterior Insula | -38 | 19 | -4 |
| Right Anterior Insula | 43 | 19 | -4 |
| Superior Frontal Gyrus | 7 | 19 | 64 |
| Left Inferior Occipital Lobe | -38 | -91 | -4 |
| Right Inferior Occipital Lobe | 34 | -91 | -1 |
| Left Middle Temporal Gyrus | -62 | -29 | -4 |
| Right Superior Temporal Gyrus | 53 | -24 | -1 |
| Right Middle Frontal Gyrus | 50 | 29 | 23 |
| Left Middle Frontal Gyrus | -44 | 26 | 20 |
| Posterior Cingulate Cortex | -2 | -31 | 28 |
| Supplementary Motor Cortex | -6 | 18 | 62 |

*Note.* Contrast = [incongruent] > [congruent]. See Figure 1 in main results.

**Table 12.**

*Peak clusters of brain regions exhibiting significant increases in activation during AX-CPT.*

| Region | x (mm) | y (mm) | z (mm) |
|---|---|---|---|
| Left Superior Parietal Lobe | -34 | -48 | 42 |
| Right Angular Gyrus | 36 | -55 | 47 |
| Right Posterior Cingulate Cortex | 2 | -31 | 25 |
| Left Inferior Occipital Lobe | -36 | -91 | -1 |
| Right Inferior Occipital Lobe | 36 | -89 | -11 |
| Left Middle Frontal Gyrus | -46 | 36 | 28 |
| Right Middle Frontal Gyrus | 48 | 38 | 28 |
| Left Anterior Insula | -31 | 17 | 8 |
| Right Anterior Insula | 34 | 19 | 6 |
| Left Precentral Gyrus | -58 | 7 | 35 |
| Left Postcentral Gyrus | -43 | -22 | 54 |
| Supplementary Motor Cortex | 2 | 14 | 47 |

*Note.* Contrast = [AX, AY, Ang, BX, BY, Bng] > [baseline]. See Figure 2 in main results.

**Table 13.**

*Peak clusters of brain regions exhibiting significant increases in activation during cued task-switching.*

| Region | x (mm) | y (mm) | z (mm) |
|---|---|---|---|
| Left Occipital Lobe | -17 | -94 | -8 |
| Right Occipital Lobe | 19 | -91 | -6 |
| Left Middle Temporal Gyrus | -55 | -41 | 4 |
| Right Middle Temporal Gyrus | 48 | -34 | 1 |
| Left Superior Temporal Gyrus | -55 | -48 | 13 |

| | | | |
|---|---|---|---|
| Right Superior Temporal Gyrus | 53 | -43 | 16 |
| Left Precentral Gyrus | -53 | 5 | 49 |
| Right Precentral Gyrus | 50 | -7 | 40 |
| Left Putamen | -24 | 5 | 11 |
| Supplementary Motor Cortex | -5 | 5 | 64 |

*Note.* Contrast = [congruent, incongruent] > [baseline]. See Figure 3 in main results.

**Table 14.**
*Peak clusters of brain regions exhibiting significant increases in activation during Sternberg task.*

| Region | x (mm) | y (mm) | z (mm) |
|---|---|---|---|
| Left Precentral Gyrus | -38 | -14 | 64 |
| Left Anterior Insula | -34 | 22 | -1 |
| Right Anterior Insula | 34 | 22 | 4 |
| Left Anterior Midcingulate Cortex | -5 | 5 | 30 |
| Right Anterior Midcingulate Cortex | 5 | 5 | 30 |
| Left Superior Parietal Lobe | -31 | -55 | 47 |
| Right Superior Parietal Lobe | 34 | -53 | 47 |
| Left Inferior Occipital Lobe | -43 | -74 | -13 |
| Right Inferior Occipital Lobe | 36 | -89 | -6 |
| Supplementary Motor Cortex | 2 | 14 | 49 |

*Note.* Contrast = [RN, NN, NP] > [baseline]. See Figure 4 in main results.

**Table 15.**
*Results of one-sample t-tests for corrected and uncorrected significance of claustrum activation across all conditions and phases during Stroop.*

| Hemisphere | Condition | P-value | FDR Corrected P-value | T-statistic |
|---|---|---|---|---|
| L | Inc | 6.416E-04 | 9.949E-04 | 3.624 |
| L | Con | 0.41 | 0.41 | 0.832 |
| R | Inc | 7.462E-04 | 9.949E-04 | 3.576 |
| R | Con | 4.049E-04 | 9.949E-04 | 3.771 |

**Table 16.**
*Results of one-sample t-tests for corrected and uncorrected significance of claustrum activation across all conditions and phases during AX-CPT.*

| Hemisphere | Condition/Phase | P-value | FDR Corrected P-value | T-statistic |
|---|---|---|---|---|
| L | AX | 2.229E-06 | 7.523E-06 | 5.296 |
| L | AX_Cue | 7.799E-06 | 1.798E-05 | 4.945 |
| L | AX_Delay | 8.827E-04 | 9.534E-04 | -3.521 |

| | | | | |
|---|---|---|---|---|
| L | AX_Probe | 1.744E-07 | 1.177E-06 | 5.994 |
| L | AY | 7.025E-07 | 2.918E-06 | 5.614 |
| L | AY_Cue | 2.153E-05 | 3.875E-05 | 4.654 |
| L | AY_Delay | 9.350E-05 | 1.365E-04 | -4.222 |
| L | AY_Probe | 1.172E-05 | 2.484E-05 | 4.829 |
| L | Ang | 4.291E-05 | 6.815E-05 | 4.453 |
| L | Ang_Cue | 1.038E-04 | 1.475E-04 | 4.190 |
| L | Ang_Delay | 2.959E-05 | 4.927E-05 | -4.562 |
| L | Ang_Probe | 3.832E-07 | 2.069E-06 | 5.780 |
| L | BX | 3.096E-06 | 8.800E-06 | 5.204 |
| L | BX_Cue | 1.826E-05 | 3.630E-05 | 4.702 |
| L | BX_Delay | 6.126E-04 | 7.191E-04 | -3.639 |
| L | BX_Probe | 6.532E-05 | 1.008E-04 | 4.329 |
| L | BY | 3.011E-05 | 4.927E-05 | 4.557 |
| L | BY_Cue | 4.777E-04 | 5.732E-04 | 3.719 |
| L | BY_Delay | 1.540E-07 | 1.177E-06 | -6.028 |
| L | BY_Probe | 1.196E-05 | 2.484E-05 | 4.823 |
| L | Bng | 7.002E-04 | 8.044E-04 | 3.596 |
| L | Bng_Cue | 8.762E-04 | 9.534E-04 | 3.523 |
| L | Bng_Delay | 2.984E-04 | 3.663E-04 | -3.867 |
| L | Bng_Probe | 2.952E-05 | 4.927E-05 | 4.562 |
| L | Cue | 2.278E-09 | 3.076E-08 | 7.158 |
| L | Delay | 2.529E-08 | 2.732E-07 | -6.514 |
| L | Probe | 7.890E-10 | 1.433E-08 | 7.442 |
| R | AX | 4.726E-06 | 1.215E-05 | 5.086 |
| R | AX_Cue | 1.259E-04 | 1.700E-04 | 4.132 |
| R | AX_Delay | 3.915E-06 | 1.057E-05 | -5.139 |

| | | | | |
|---|---|---|---|---|
| R | AX_Probe | 9.701E-08 | 8.731E-07 | 6.153 |
| R | AY | 8.149E-04 | 9.167E-04 | 3.547 |
| R | AY_Cue | 3.839E-02 | 0.038 | 2.123 |
| R | AY_Delay | 3.223E-03 | 3.347E-03 | -3.083 |
| R | AY_Probe | 1.976E-05 | 3.680E-05 | 4.679 |
| R | Ang | 5.466E-06 | 1.342E-05 | 5.045 |
| R | Ang_Cue | 2.402E-04 | 3.088E-04 | 3.934 |
| R | Ang_Delay | 5.774E-07 | 2.598E-06 | -5.668 |
| R | Ang_Probe | 5.232E-07 | 2.568E-06 | 5.695 |
| R | BX | 2.627E-06 | 8.247E-06 | 5.250 |
| R | BX_Cue | 9.267E-05 | 1.365E-04 | 4.224 |
| R | BX_Delay | 2.156E-06 | 7.523E-06 | -5.305 |
| R | BX_Probe | 1.001E-06 | 3.860E-06 | 5.517 |
| R | BY | 1.085E-04 | 1.503E-04 | 4.177 |
| R | BY_Cue | 1.097E-03 | 1.161E-03 | 3.450 |
| R | BY_Delay | 7.991E-06 | 1.798E-05 | -4.938 |
| R | BY_Probe | 2.749E-06 | 8.247E-06 | 5.237 |
| R | Bng | 2.935E-04 | 3.663E-04 | 3.872 |
| R | Bng_Cue | 3.605E-03 | 3.673E-03 | 3.044 |
| R | Bng_Delay | 2.071E-04 | 2.728E-04 | -3.980 |
| R | Bng_Probe | 1.882E-05 | 3.630E-05 | 4.693 |
| R | Cue | 2.134E-07 | 1.280E-06 | 5.939 |
| R | Delay | 3.123E-10 | 1.433E-08 | -7.690 |
| R | Probe | 7.963E-10 | 1.433E-08 | 7.440 |

**Table 17.**

*Results of one-sample t-tests for corrected and uncorrected significance of claustrum activation across all trial types and phases during cued task-switching.*

| Hemisphere | Condition/Phase | P-value | FDR Corrected P-value | T-statistic |
|---|---|---|---|---|
| L | Con | 0.230 | 0.321 | 1.215 |
| L | Con_Cue | 0.469 | 0.525 | -0.729 |
| L | Con_Delay | 1.405E-04 | 0.001 | 4.099 |
| L | Con_Feedback | 0.394 | 0.460 | 0.858 |
| L | Con_Probe | 0.037 | 0.097 | 2.140 |
| L | Inc | 0.052 | 0.117 | 1.985 |
| L | Inc_Cue | 0.899 | 0.899 | -0.128 |
| L | Inc_Delay | 0.001 | 0.005 | 3.476 |
| L | Inc_Feedback | 0.035 | 0.097 | 2.157 |
| L | Inc_Probe | 0.062 | 0.117 | 1.907 |
| L | Cue | 0.647 | 0.671 | -0.461 |
| L | Delay | 5.042E-05 | 7.06E-04 | 4.406 |
| L | Feedback | 0.071 | 0.117 | 1.840 |
| L | Probe | 0.019 | 0.065 | 2.428 |
| R | Con | 0.310 | 0.377 | -1.025 |
| R | Con_Cue | 0.490 | 0.528 | -0.695 |
| R | Con_Delay | 1.490E-04 | 0.001 | 4.081 |
| R | Con_Feedback | 0.016 | 0.064 | -2.485 |
| R | Con_Probe | 0.068 | 0.117 | 1.866 |
| R | Inc | 0.294 | 0.374 | -1.060 |
| R | Inc_Cue | 0.131 | 0.203 | -1.535 |
| R | Inc_Delay | 2.959E-04 | 0.002 | 3.870 |
| R | Inc_Feedback | 0.289 | 0.374 | -1.070 |
| R | Inc_Probe | 0.067 | 0.117 | 1.867 |

| | | | | |
|---|---|---|---|---|
| R | Cue | 0.224 | 0.321 | -1.230 |
| R | Delay | 4.461E-05 | 7.06E-04 | 4.442 |
| R | Feedback | 0.068 | 0.117 | -1.863 |
| R | Probe | 0.038 | 0.097 | 2.124 |

**Table 18.**

*Results of one-sample t-tests for corrected and uncorrected significance across all trial switches and phases during cued task-switching.*

| Hemisphere | Condition/Phase | P-value | FDR Corrected P-value | T-statistic |
|---|---|---|---|---|
| L | Repeat | 0.214 | 0.329 | 1.258 |
| L | Repeat_Cue | 0.685 | 0.685 | -0.408 |
| L | Repeat_Delay | 0.001 | 0.005 | 3.491 |
| L | Repeat_Feedback | 0.468 | 0.502 | 0.731 |
| L | Repeat_Probe | 0.018 | 0.074 | 2.430 |
| L | Switch | 0.154 | 0.257 | 1.445 |
| L | Switch_Cue | 0.445 | 0.502 | -0.770 |
| L | Switch_Delay | 6.56E-05 | 9.45E-04 | 4.328 |
| L | Switch_Feedback | 0.072 | 0.160 | 1.835 |
| L | Switch_Probe | 0.080 | 0.160 | 1.786 |
| R | Repeat | 0.370 | 0.463 | -0.904 |
| R | Repeat_Cue | 0.477 | 0.502 | -0.716 |
| R | Repeat_Delay | 0.001 | 0.004 | 3.676 |
| R | Repeat_Feedback | 0.055 | 0.158 | -1.958 |
| R | Repeat_Probe | 0.047 | 0.156 | 2.035 |
| R | Switch | 0.140 | 0.255 | -1.496 |

| | | | | |
|---|---|---|---|---|
| R | Switch_Cue | 0.075 | 0.160 | -1.814 |
| R | Switch_Delay | 9.45E-05 | 9.45E-04 | 4.219 |
| R | Switch_Feedback | 0.270 | 0.385 | -1.116 |
| R | Switch_Probe | 0.312 | 0.416 | 1.020 |

**Table 19.**

*Results of one-sample t-tests for corrected and uncorrected significance across all conditions and phases during Sternberg.*

| Hemisphere | Condition/Phase | P-value | FDR Corrected P-value | T-statistic |
|---|---|---|---|---|
| L | NN | 1.578E-08 | 7.891E-08 | 6.641 |
| L | NN_Encoding | 7.044E-02 | 0.101 | 1.846 |
| L | NN_Retention | 2.972E-01 | 0.388 | 1.053 |
| L | NN_Retrieval | 8.723E-09 | 6.543E-08 | 6.799 |
| L | NP | 1.153E-08 | 6.918E-08 | 6.725 |
| L | NP_Encoding | 5.973E-03 | 9.955E-03 | 2.862 |
| L | NP_Retention | 8.555E-01 | 0.885 | 0.183 |
| L | NP_Retrieval | 2.536E-09 | 2.536E-08 | 7.130 |
| L | RN | 9.158E-07 | 3.053E-06 | 5.542 |
| L | RN_Encoding | 2.933E-04 | 5.866E-04 | 3.872 |
| L | RN_Retention | 4.471E-01 | 0.497 | -0.766 |
| L | RN_Retrieval | 4.824E-05 | 1.034E-04 | 4.419 |
| L | Encoding | 9.714E-04 | 1.714E-03 | 3.490 |
| L | Retention | 9.925E-01 | 0.993 | -0.009 |
| L | Retrieval | 8.444E-10 | 1.267E-08 | 7.424 |
| R | NN | 7.765E-06 | 2.118E-05 | 4.946 |
| R | NN_Encoding | 3.166E-01 | 0.396 | -1.011 |

| | | | | |
|---|---|---|---|---|
| R | NN_Retention | 2.044E-02 | 3.066E-02 | 2.389 |
| R | NN_Retrieval | 2.085E-06 | 6.256E-06 | 5.314 |
| R | NP | 1.217E-05 | 3.042E-05 | 4.818 |
| R | NP_Encoding | 4.133E-01 | 0.477 | -0.824 |
| R | NP_Retention | 3.398E-04 | 6.372E-04 | 3.826 |
| R | NP_Retrieval | 1.722E-05 | 3.974E-05 | 4.718 |
| R | RN | 1.967E-08 | 8.431E-08 | 6.582 |
| R | RN_Encoding | 5.874E-01 | 0.629 | -0.546 |
| R | RN_Retention | 2.727E-01 | 0.372 | 1.108 |
| R | RN_Retrieval | 3.148E-07 | 1.181E-06 | 5.834 |
| R | Encoding | 3.487E-01 | 0.419 | -0.945 |
| R | Retention | 9.307E-03 | 0.015 | 2.697 |
| R | Retrieval | 5.767E-10 | 1.267E-08 | 7.526 |

**Table 20.**

*Variance inflation factors by phase and session for each subject during AX-CPT.*

| Subject | Ses 1 Cue | Ses 1 Delay | Ses 1 Probe | Ses 2 Cue | Ses 2 Delay | Ses 2 Probe |
|---|---|---|---|---|---|---|
| sub-f1027ao | 1.617 | 2.062 | 1.536 | 1.615 | 2.044 | 1.536 |
| sub-f1031ax | 1.618 | 2.039 | 1.537 | 1.613 | 2.064 | 1.532 |
| sub-f1342ku | 1.617 | 2.053 | 1.536 | 1.615 | 2.030 | 1.535 |
| sub-f1550bc | 1.619 | 2.030 | 1.540 | 1.620 | 2.054 | 1.540 |
| sub-f1552xo | 1.617 | 2.065 | 1.538 | 1.613 | 2.042 | 1.534 |
| sub-f1659oa | 1.622 | 2.052 | 1.542 | 1.615 | 2.056 | 1.534 |
| sub-f1670rz | 1.631 | 2.081 | 1.548 | 1.617 | 2.034 | 1.538 |
| sub-f1828ko | 1.611 | 2.068 | 1.530 | 1.617 | 2.034 | 1.538 |
| sub-f2157me | 1.625 | 2.043 | 1.546 | 1.620 | 2.047 | 1.540 |

| | | | | | | |
|---|---|---|---|---|---|---|
| sub-f2499cq | 1.609 | 2.077 | 1.529 | 1.615 | 2.067 | 1.534 |
| sub-f2593wi | 1.613 | 2.051 | 1.532 | 1.618 | 2.018 | 1.538 |
| sub-f2648qw | 1.613 | 2.014 | 1.533 | 1.617 | 2.036 | 1.536 |
| sub-f2709ul | 1.622 | 2.052 | 1.540 | 1.612 | 2.042 | 1.533 |
| sub-f2968sp | 1.618 | 2.042 | 1.537 | 1.612 | 2.030 | 1.532 |
| sub-f3300jh | 1.615 | 2.030 | 1.536 | 1.612 | 2.033 | 1.531 |
| sub-f3387yq | 1.615 | 2.055 | 1.535 | 1.613 | 2.049 | 1.533 |
| sub-f3469wa | 1.611 | 2.068 | 1.532 | 1.628 | 2.087 | 1.546 |
| sub-f3526dz | 1.625 | 2.090 | 1.545 | 1.614 | 2.029 | 1.534 |
| sub-f3680fb | 1.616 | 2.065 | 1.535 | 1.617 | 2.045 | 1.537 |
| sub-f3681wf | 1.613 | 2.046 | 1.533 | 1.615 | 2.044 | 1.536 |
| sub-f3996sp | 1.612 | 2.035 | 1.533 | 1.610 | 2.028 | 1.532 |
| sub-f4138ge | 1.619 | 2.033 | 1.539 | 1.618 | 2.026 | 1.539 |
| sub-f4310gw | 1.620 | 2.057 | 1.539 | 1.610 | 2.026 | 1.531 |
| sub-f4354bs | 1.622 | 2.071 | 1.542 | 1.616 | 2.019 | 1.537 |
| sub-f4467ur | 1.614 | 2.037 | 1.535 | 1.625 | 2.076 | 1.543 |
| sub-f4796rs | 1.621 | 2.046 | 1.540 | 1.619 | 2.063 | 1.539 |
| sub-f4831tn | 1.623 | 2.076 | 1.543 | 1.628 | 2.081 | 1.547 |
| sub-f5001ob | 1.621 | 2.061 | 1.541 | 1.619 | 2.062 | 1.539 |
| sub-f5004cr | 1.617 | 2.060 | 1.536 | 1.613 | 2.046 | 1.532 |
| sub-f5094na | 1.610 | 2.039 | 1.530 | 1.614 | 2.026 | 1.533 |
| sub-f5094ya | 1.619 | 2.068 | 1.540 | 1.611 | 2.023 | 1.532 |
| sub-f5386yx | 1.621 | 2.042 | 1.542 | 1.623 | 2.040 | 1.542 |
| sub-f5416zj | 1.622 | 2.054 | 1.543 | 1.618 | 2.060 | 1.539 |
| sub-f5445nh | 1.620 | 2.056 | 1.541 | 1.617 | 2.078 | 1.538 |
| sub-f5635rv | 1.619 | 2.082 | 1.539 | 1.614 | 2.055 | 1.533 |
| sub-f5650zm | 1.619 | 2.071 | 1.540 | 1.620 | 2.072 | 1.540 |
| sub-f5930vp | 1.619 | 2.055 | 1.539 | 1.620 | 2.057 | 1.538 |
| sub-f6188io | 1.616 | 2.056 | 1.536 | 1.622 | 2.018 | 1.541 |

| | | | | | | |
|---|---|---|---|---|---|---|
| sub-f6318if | 1.614 | 2.069 | 1.534 | 1.624 | 2.070 | 1.543 |
| sub-f6464bf | 1.618 | 2.085 | 1.536 | 1.621 | 2.027 | 1.540 |
| sub-f6950qp | 1.621 | 2.072 | 1.539 | 1.620 | 2.070 | 1.540 |
| sub-f7227ag | 1.613 | 2.034 | 1.532 | 1.617 | 2.058 | 1.536 |
| sub-f7688lh | 1.616 | 2.075 | 1.537 | 1.622 | 2.046 | 1.542 |
| sub-f7951pz | 1.620 | 2.067 | 1.540 | 1.617 | 2.058 | 1.537 |
| sub-f8113do | 1.617 | 2.041 | 1.536 | 1.619 | 2.074 | 1.539 |
| sub-f8194sp | 1.615 | 2.050 | 1.534 | 1.611 | 2.064 | 1.532 |
| sub-f8270up | 1.612 | 2.060 | 1.531 | 1.627 | 2.085 | 1.546 |
| sub-f8294bu | 1.613 | 2.082 | 1.532 | 1.616 | 2.045 | 1.537 |
| sub-f8298ds | 1.621 | 2.030 | 1.539 | 1.612 | 2.016 | 1.533 |
| sub-f8570ui | 1.625 | 2.047 | 1.545 | 1.616 | 2.061 | 1.537 |
| sub-f8710qa | 1.626 | 2.075 | 1.544 | 1.622 | 2.069 | 1.540 |
| sub-f8979ai | 1.607 | 2.032 | 1.529 | 1.610 | 2.045 | 1.530 |
| sub-f9057kp | 1.618 | 2.036 | 1.537 | 1.612 | 2.071 | 1.531 |
| sub-f9206gd | 1.613 | 2.029 | 1.534 | 1.623 | 2.059 | 1.543 |
| sub-f9271ex | 1.622 | 2.050 | 1.543 | 1.620 | 2.060 | 1.539 |

**Table 21.**

*Variance inflation factors by phase and session for each subject during Sternberg.*

| Subject | Ses 1 Encoding | Ses 1 Retention | Ses 1 Retrieval | Ses 2 Encoding | Ses 2 Retention | Ses 2 Retrieval |
|---|---|---|---|---|---|---|
| sub-f1027ao | 1.441 | 1.132 | 1.581 | 1.427 | 1.131 | 1.565 |
| sub-f1031ax | 1.418 | 1.133 | 1.561 | 1.442 | 1.134 | 1.586 |
| sub-f1342ku | 1.475 | 1.133 | 1.617 | 1.422 | 1.132 | 1.561 |
| sub-f1550bc | 1.448 | 1.133 | 1.589 | 1.463 | 1.131 | 1.602 |
| sub-f1552xo | 1.434 | 1.133 | 1.575 | 1.418 | 1.133 | 1.561 |
| sub-f1659oa | 1.444 | 1.131 | 1.583 | 1.433 | 1.133 | 1.575 |
| sub-f1670rz | 1.442 | 1.134 | 1.585 | 1.409 | 1.132 | 1.548 |

| | | | | | | |
|---|---|---|---|---|---|---|
| sub-f1828ko | 1.440 | 1.132 | 1.580 | 1.465 | 1.132 | 1.606 |
| sub-f2157me | 1.436 | 1.133 | 1.578 | 1.427 | 1.132 | 1.566 |
| sub-f2499cq | 1.419 | 1.131 | 1.558 | 1.439 | 1.132 | 1.581 |
| sub-f2593wi | 1.445 | 1.131 | 1.583 | 1.451 | 1.133 | 1.593 |
| sub-f2648qw | 1.466 | 1.132 | 1.608 | 1.407 | 1.132 | 1.545 |
| sub-f2709ul | 1.423 | 1.134 | 1.566 | 1.445 | 1.133 | 1.587 |
| sub-f2968sp | 1.430 | 1.134 | 1.574 | 1.457 | 1.131 | 1.595 |
| sub-f3300jh | 1.443 | 1.133 | 1.585 | 1.419 | 1.133 | 1.561 |
| sub-f3387yq | 1.458 | 1.131 | 1.596 | 1.446 | 1.133 | 1.589 |
| sub-f3469wa | 1.413 | 1.132 | 1.551 | 1.457 | 1.133 | 1.599 |
| sub-f3526dz | 1.411 | 1.132 | 1.548 | 1.440 | 1.132 | 1.581 |
| sub-f3680fb | 1.432 | 1.133 | 1.572 | 1.443 | 1.133 | 1.585 |
| sub-f3681wf | 1.460 | 1.131 | 1.597 | 1.459 | 1.131 | 1.598 |
| sub-f3996sp | 1.426 | 1.134 | 1.568 | 1.400 | 1.132 | 1.539 |
| sub-f4138ge | 1.432 | 1.133 | 1.574 | 1.435 | 1.133 | 1.576 |
| sub-f4310gw | 1.427 | 1.134 | 1.570 | 1.472 | 1.133 | 1.614 |
| sub-f4354bs | 1.427 | 1.133 | 1.568 | 1.444 | 1.132 | 1.585 |
| sub-f4467ur | 1.431 | 1.134 | 1.574 | 1.457 | 1.133 | 1.599 |
| sub-f4796rs | 1.431 | 1.133 | 1.572 | 1.425 | 1.134 | 1.569 |
| sub-f4831tn | 1.424 | 1.133 | 1.565 | 1.449 | 1.133 | 1.590 |
| sub-f5001ob | 1.447 | 1.133 | 1.590 | 1.399 | 1.132 | 1.537 |
| sub-f5004cr | 1.445 | 1.133 | 1.585 | 1.431 | 1.133 | 1.573 |
| sub-f5094na | 1.463 | 1.132 | 1.603 | 1.478 | 1.133 | 1.620 |
| sub-f5094ya | 1.470 | 1.132 | 1.611 | 1.430 | 1.131 | 1.567 |
| sub-f5386yx | 1.428 | 1.134 | 1.571 | 1.460 | 1.133 | 1.601 |

| | | | | | | |
|---|---|---|---|---|---|---|
| sub-f5416zj | 1.438 | 1.134 | 1.581 | 1.440 | 1.134 | 1.584 |
| sub-f5445nh | 1.410 | 1.132 | 1.549 | 1.437 | 1.131 | 1.575 |
| sub-f5635rv | 1.401 | 1.132 | 1.538 | 1.402 | 1.132 | 1.542 |
| sub-f5650zm | 1.439 | 1.131 | 1.577 | 1.441 | 1.133 | 1.581 |
| sub-f5930vp | 1.406 | 1.132 | 1.545 | 1.447 | 1.133 | 1.590 |
| sub-f6188io | 1.475 | 1.133 | 1.617 | 1.413 | 1.133 | 1.554 |
| sub-f6318if | 1.447 | 1.133 | 1.589 | 1.430 | 1.134 | 1.572 |
| sub-f6464bf | 1.451 | 1.133 | 1.593 | 1.403 | 1.131 | 1.540 |
| sub-f6950qp | 1.438 | 1.134 | 1.581 | 1.436 | 1.132 | 1.576 |
| sub-f7227ag | 1.472 | 1.133 | 1.614 | 1.450 | 1.133 | 1.591 |
| sub-f7688lh | 1.448 | 1.133 | 1.590 | 1.418 | 1.133 | 1.558 |
| sub-f7951pz | 1.428 | 1.132 | 1.566 | 1.420 | 1.133 | 1.561 |
| sub-f8113do | 1.428 | 1.134 | 1.571 | 1.446 | 1.133 | 1.588 |
| sub-f8194sp | 1.412 | 1.133 | 1.553 | 1.454 | 1.134 | 1.598 |
| sub-f8270up | 1.431 | 1.131 | 1.568 | 1.412 | 1.133 | 1.552 |
| sub-f8294bu | 1.448 | 1.133 | 1.590 | 1.431 | 1.133 | 1.574 |
| sub-f8298ds | 1.458 | 1.133 | 1.601 | 1.460 | 1.133 | 1.604 |
| sub-f8570ui | 1.432 | 1.133 | 1.574 | 1.459 | 1.133 | 1.603 |
| sub-f8710qa | 1.437 | 1.132 | 1.577 | 1.448 | 1.133 | 1.590 |
| sub-f8979ai | 1.399 | 1.132 | 1.536 | 1.456 | 1.134 | 1.598 |
| sub-f9057kp | 1.419 | 1.134 | 1.561 | 1.425 | 1.132 | 1.564 |
| sub-f9206gd | 1.428 | 1.133 | 1.569 | 1.455 | 1.134 | 1.599 |
| sub-f9271ex | 1.441 | 1.134 | 1.585 | 1.432 | 1.131 | 1.571 |

**Table 22.**

*Variance inflation factors by phase and session for each subject during cued task-switching.*

| Subject | Ses 1 Cue | Ses 1 Delay | Ses 1 Probe | Ses1 Feedback | Ses 2 Cue | Ses 2 Delay | Ses 2 Probe | Ses 2 Feedback |
|---|---|---|---|---|---|---|---|---|
| sub-f1027ao | 3.335 | 3.846 | 4.580 | 3.778 | 3.338 | 3.832 | 4.535 | 3.757 |
| sub-f1031ax | 3.324 | 3.840 | 4.561 | 3.760 | 3.331 | 3.832 | 4.566 | 3.765 |
| sub-f1342ku | 3.343 | 3.838 | 4.571 | 3.779 | 3.332 | 3.831 | 4.568 | 3.765 |
| sub-f1550bc | 3.331 | 3.847 | 4.571 | 3.771 | 3.318 | 3.829 | 4.543 | 3.745 |
| sub-f1552xo | 3.341 | 3.834 | 4.569 | 3.779 | 3.337 | 3.842 | 4.574 | 3.779 |
| sub-f1659oa | 3.316 | 3.831 | 4.560 | 3.761 | 3.330 | 3.848 | 4.580 | 3.776 |
| sub-f1670rz | 3.330 | 3.847 | 4.585 | 3.776 | 3.326 | 3.826 | 4.546 | 3.750 |
| sub-f1828ko | 3.318 | 3.823 | 4.518 | 3.729 | 3.333 | 3.833 | 4.581 | 3.786 |
| sub-f2157me | 3.325 | 3.827 | 4.543 | 3.753 | 3.344 | 3.840 | 4.575 | 3.781 |
| sub-f2499cq | 3.342 | 3.844 | 4.558 | 3.764 | 3.340 | 3.834 | 4.561 | 3.770 |
| sub-f2593wi | 3.329 | 3.844 | 4.576 | 3.773 | 3.331 | 3.839 | 4.588 | 3.783 |
| sub-f2648qw | 3.347 | 3.832 | 4.550 | 3.765 | 3.340 | 3.832 | 4.563 | 3.772 |
| sub-f2709ul | 3.326 | 3.826 | 4.543 | 3.758 | 3.328 | 3.831 | 4.568 | 3.767 |
| sub-f2968sp | 3.329 | 3.828 | 4.566 | 3.774 | 3.363 | 3.852 | 4.583 | 3.797 |
| sub-f3300jh | 3.345 | 3.843 | 4.575 | 3.784 | 3.354 | 3.839 | 4.547 | 3.771 |
| sub-f3387yq | 3.345 | 3.842 | 4.573 | 3.785 | 3.313 | 3.823 | 4.546 | 3.750 |
| sub-f3469wa | 3.337 | 3.836 | 4.566 | 3.776 | 3.349 | 3.847 | 4.581 | 3.795 |
| sub-f3526dz | 3.330 | 3.843 | 4.574 | 3.768 | 3.342 | 3.834 | 4.570 | 3.779 |
| sub-f3680fb | 3.318 | 3.830 | 4.560 | 3.762 | 3.322 | 3.830 | 4.564 | 3.768 |
| sub-f3681wf | 3.327 | 3.831 | 4.557 | 3.760 | 3.337 | 3.830 | 4.533 | 3.752 |
| sub-f3996sp | 3.329 | 3.830 | 4.569 | 3.772 | 3.347 | 3.841 | 4.564 | 3.780 |
| sub-f4138ge | 3.329 | 3.826 | 4.544 | 3.760 | 3.312 | 3.822 | 4.533 | 3.740 |
| sub-f4310gw | 3.341 | 3.842 | 4.582 | 3.792 | 3.332 | 3.832 | 4.541 | 3.751 |

| | | | | | | | | |
|---|---|---|---|---|---|---|---|---|
| sub-f4354bs | 3.334 | 3.845 | 4.577 | 3.772 | 3.330 | 3.833 | 4.573 | 3.774 |
| sub-f4467ur | 3.332 | 3.843 | 4.565 | 3.762 | 3.346 | 3.845 | 4.589 | 3.792 |
| sub-f4796rs | 3.315 | 3.833 | 4.567 | 3.759 | 3.336 | 3.841 | 4.560 | 3.762 |
| sub-f4831tn | 3.343 | 3.843 | 4.562 | 3.767 | 3.341 | 3.840 | 4.571 | 3.781 |
| sub-f5001ob | 3.330 | 3.827 | 4.562 | 3.763 | 3.317 | 3.826 | 4.558 | 3.757 |
| sub-f5004cr | 3.329 | 3.830 | 4.523 | 3.742 | 3.335 | 3.832 | 4.575 | 3.778 |
| sub-f5094na | 3.318 | 3.834 | 4.549 | 3.742 | 3.337 | 3.837 | 4.570 | 3.774 |
| sub-f5094ya | 3.316 | 3.830 | 4.559 | 3.752 | 3.320 | 3.835 | 4.569 | 3.758 |
| sub-f5386yx | 3.332 | 3.838 | 4.576 | 3.778 | 3.323 | 3.832 | 4.563 | 3.758 |
| sub-f5416zj | 3.333 | 3.843 | 4.573 | 3.771 | 3.348 | 3.856 | 4.606 | 3.798 |
| sub-f5445nh | 3.336 | 3.827 | 4.538 | 3.750 | 3.318 | 3.828 | 4.551 | 3.750 |
| sub-f5635rv | 3.318 | 3.830 | 4.560 | 3.763 | 3.350 | 3.848 | 4.576 | 3.778 |
| sub-f5650zm | 3.313 | 3.835 | 4.545 | 3.741 | 3.317 | 3.824 | 4.560 | 3.759 |
| sub-f5930vp | 3.335 | 3.838 | 4.582 | 3.788 | 3.313 | 3.825 | 4.544 | 3.742 |
| sub-f6188io | 3.319 | 3.830 | 4.561 | 3.763 | 3.332 | 3.834 | 4.566 | 3.773 |
| sub-f6318if | 3.310 | 3.826 | 4.560 | 3.758 | 3.314 | 3.821 | 4.549 | 3.748 |
| sub-f6464bf | 3.312 | 3.827 | 4.538 | 3.745 | 3.320 | 3.829 | 4.565 | 3.756 |
| sub-f6950qp | 3.322 | 3.828 | 4.552 | 3.755 | 3.323 | 3.821 | 4.535 | 3.743 |
| sub-f7227ag | 3.358 | 3.848 | 4.585 | 3.800 | 3.340 | 3.839 | 4.568 | 3.778 |
| sub-f7688lh | 3.325 | 3.829 | 4.554 | 3.762 | 3.323 | 3.833 | 4.573 | 3.775 |
| sub-f7951pz | 3.323 | 3.828 | 4.541 | 3.756 | 3.348 | 3.854 | 4.591 | 3.790 |
| sub-f8113do | 3.330 | 3.832 | 4.574 | 3.777 | 3.334 | 3.835 | 4.563 | 3.771 |
| sub-f8194sp | 3.343 | 3.841 | 4.552 | 3.771 | 3.339 | 3.827 | 4.537 | 3.754 |
| sub-f8270up | 3.317 | 3.837 | 4.581 | 3.768 | 3.314 | 3.832 | 4.566 | 3.757 |
| sub-f8294bu | 3.321 | 3.830 | 4.567 | 3.769 | 3.329 | 3.842 | 4.568 | 3.768 |

| | | | | | | | | |
|---|---|---|---|---|---|---|---|---|
| sub-f8298ds | 3.314 | 3.831 | 4.546 | 3.745 | 3.312 | 3.829 | 4.564 | 3.764 |
| sub-f8570ui | 3.316 | 3.833 | 4.564 | 3.758 | 3.323 | 3.839 | 4.571 | 3.767 |
| sub-f8710qa | 3.331 | 3.843 | 4.580 | 3.778 | 3.330 | 3.849 | 4.578 | 3.774 |
| sub-f8979ai | 3.314 | 3.829 | 4.562 | 3.763 | 3.333 | 3.831 | 4.553 | 3.762 |
| sub-f9057kp | 3.329 | 3.840 | 4.576 | 3.769 | 3.350 | 3.852 | 4.595 | 3.802 |
| sub-f9206gd | 3.332 | 3.840 | 4.567 | 3.766 | 3.319 | 3.829 | 4.556 | 3.761 |
| sub-f9271ex | 3.344 | 3.842 | 4.590 | 3.793 | 3.318 | 3.832 | 4.566 | 3.756 |
